# Depletion of neurocan in the prefrontal cortex impairs temporal order recognition, cognitive flexibility and perisomatic GABAergic innervation

**DOI:** 10.1101/2023.04.18.537277

**Authors:** David Baidoe-Ansah, Hadi Mirzapourdelavar, Hector Carceller, Marta Pérez-Rando, Luisa Strackeljan, Borja Garcia-Vazquez, Constanze Seidenbecher, Rahul Kaushik, Juan Nacher, Alexander Dityatev

## Abstract

The condensed form of neural extracellular matrix (ECM), perineuronal nets (PNNs), is predominantly associated with parvalbumin-expressing (PV+) interneurons in the cortex and hippocampus. PNNs are enriched in several lecticans, including neurocan (Ncan). A polymorphism in the human *Ncan* gene has been associated with alterations in hippocampus-dependent memory function, variation of prefrontal cortex structure, and a higher risk for schizophrenia or bipolar disorder. Ncan knockout (KO) mice show related behavioral abnormalities, such as hyperactivity. Here we focused on studying how dysregulation of Ncan specifically in the mPFC may affect cognitive and synaptic functions. Intracortical adeno-associated virus (AAV) delivery was used to express shRNA against Ncan. Analysis of PNNs in Ncan shRNA-injected mice revealed a reduction in PNNs labelling by *Wisteria floribunda* agglutinin (WFA) around PV+ interneurons. Reduced Ncan expression resulted in a loss of the mPFC-dependent temporal order recognition and impairment of reversal spatial learning in a labyrinth (dry maze) task. As a potential synaptic substrate of these cognitive abnormalities, we report a robust reduction in the perisomatic GABAergic innervation of PV+ cells in Ncan KO and Ncan shRNA-injected mice. We also observed an increase in the density of vGLUT1-immunopositive synaptic puncta in the neuropil of Ncan shRNA-injected mice, which was, however, compensated in Ncan KO mice. Thus, our findings highlight a functional role of Ncan in supporting perisomatic GABAergic inhibition, temporal order recognition memory and cognitive flexibility, as one of the important cognitive resources depleted in neuropsychiatric disorders.

**Contribution to the field:** In this study, we asked if the extracellular matrix proteoglycan neurocan (Ncan) plays a functional role in the prefrontal cortex (PFC) of mice. Using viral delivery and expression of shRNA to knock down the expression of *Ncan* in the PFC, we provide evidence that neuronal Ncan is essential for the maintenance of perineuronal nets enveloping perisomatic interneurons by influencing the expression of glycoepitopes stained with *Wisteria floribunda* agglutinin and by modulating mRNA expression levels of other PNNs constituents. At the behavioral level, the knockdown of Ncan in mPFC impaired the temporal order recognition memory and consolidation/retrieval of spatial memories after reversal learning in the dry maze task. At the synaptic level, we found that Ncan knockdown reduced perisomatic GABAergic innervation of perisomatic interneurons and increased the density of vGLUT1+ excitatory presynaptic terminals in the neuropil of the PFC. Moreover, knockdown of Ncan changed the expression levels of several genes involved in activity-dependent synaptic remodeling. In summary, we conclude that neuronal Ncan is essential for multiple cognitive flexibility-related synaptic and cognitive functions in the PFC.

## INTRODUCTION

The medial prefrontal cortex (mPFC) has been identified as a critical domain for long-term memory storage (Euston et al., 2012; Hylin et al., 2013) and decision-making (Rushworth et al., 2011; Domenech, 2015; Funahashi, 2017; Domenech et al., 2020). Additionally, the mPFC has been shown to contribute significantly to the retrieval phase of memory remodeling (Peters et al., 2013; Hauser et al., 2015), and during cognitive flexibility (CF) (Coutlee and Huettel, 2012; Hauser et al., 2015). The latter is a component of the executive function, which includes task-shifting properties (Hauser et al., 2015). It is defined as the ability to adapt to a steady dynamic environment as well as to filter information to focus on relevant features in an impending task (Monsell, 2003; Happel and Frischknecht, 2016; Buttelmann and Karbach, 2017). CF is an essential component during reversal learning paradigms (Izquierdo et al., 2017), which have been shown to depend on the integrity of the extracellular matrix (ECM) (Morellini et al., 2010; Happel et al., 2014).

The interstitial space of the brain is known to encompass diverse structural and functional families of ECM molecules, including tenascins, heparan sulfate proteoglycans (HSPGs) and chondroitin sulfate proteoglycans (CSPGs). CSPGs are also incorporated in the condensed form of neural ECM, known as perineuronal nets (PNNs), which mostly surround soma, proximal dendrites, and axon initial segments of parvalbumin-expressing (PV+) interneurons (Dityatev and Schachner, 2003; Bruckner et al., 2006; Fawcett et al., 2019). PNNs are formed in an activity-dependent manner (Dityatev et al., 2007) and are broadly detectable in the hippocampus, mPFC, cerebellum and other brain areas (Morris and Henderson, 2000; Bruckner et al., 2006). PNN formation and maturation coincide with the onset and closure of the critical period that can be reopened by PNN disintegration (Pizzorusso et al., 2002; Hou et al., 2017). Functionally, PNNs have been associated with neural processes such as synaptic plasticity as well as formation, stabilization, updating and recall of memory (Dityatev and Fellin, 2008; Gogolla et al., 2009; Happel and Frischknecht, 2016; Thompson and Chen, 2017). Interestingly, the presence of PNNs is an important regulator of the synaptic input to PV+ interneurons, particularly of their GABAergic innervation (Carceller et al., 2020).

PNNs are composed of lecticans, namely aggrecan (Acan), versican (Vcan), brevican (Bcan), and neurocan (Ncan), which all bind to hyaluronan as a backbone and are stabilized by hyaluronan and proteoglycan link proteins (HAPLN1-4), and cross-linked by the glycoprotein tenascin-R (TnR) (Dityatev et al., 2010; Morawski et al., 2014). A study by Suttkus and colleagues dissected the importance of individual ECM molecules in the somatosensory cortex (Suttkus et al., 2014) using KO mice, and identified Acan, HAPLN1, and TnR, but not Bcan, as essential contributors to PNN formation (Suttkus et al., 2014). Their findings also correlate with other studies (Yamada et al., 2017). Interestingly, a KO study in the medial nucleus of the trapezoid body (MNTB) of mice implicated Ncan in PNNs development, potentially via modulation of the expression pattern of other PNN molecules such as HAPLN1 and Bcan (Schmidt et al., 2020).

Ncan, a brain-specific CSPG, is known to affect neuronal adhesion and migration by interacting with the neural cell adhesion molecule NCAM (Raum et al., 2015). Expression of Ncan is postnatally downregulated and later upregulated again during adulthood (Pizzorusso et al., 2002). This pattern of expression correlates with the inhibitory phenotype of mature CSPGs toward axonal growth, which is reversed once CSPGs are digested (Pizzorusso et al., 2002). Similarly, Ncan is upregulated during glial scar formation after injury, where it prevents axonal growth (Asher et al., 2000).

The human *Ncan* gene single-nucleotide polymorphism (SNP) rs1064395 is a well-documented risk factor for schizophrenia and bipolar disorder. Miró and colleagues found an association between this SNP and some behavioral abnormalities such as mania factor (hyperactivity) in humans and Ncan KO mice (Miro et al., 2012). In the pentylenetetrazole-induced model of epilepsy, Ncan expression in the mPFC has been shown to decrease along with Acan and Tn-R (Chen et al., 2016). Importantly, the mentioned risk variant in the Ncan gene (rs1064395) is associated with reduced hippocampus-dependent memory function, and variation of prefrontal cortex structure and ECM composition in healthy humans (Assmann et al., 2020), and in bipolar disorder patients (Cichon et al., 2011).

Several previous studies have used *Ncan* KO mice (Zhou et al., 2001; Schmidt et al., 2020) and enzymatic ECM digestion (Shi et al., 2019) to dissect the role of Ncan, however, these approaches have some limitations. For example, there can be activation of compensatory mechanisms, particularly during early development, in KO models (Morawski et al., 2014; El-Brolosy and Stainier, 2017) while the global acute digestion of ECM with enzymes such as chondroitinase ABC (chABC) or hyaluronidase (Dityatev et al., 2007; Happel et al., 2014; Yang et al., 2015) affects multiple ECM molecules. Thus, in this study, we employed a short hairpin RNA (shRNA)-mediated knockdown approach to study the functions of Ncan in the adult mPFC. This technique is molecular target-specific and can be applied after cessation of major developmental events in a specific brain region (O’Keefe, 2013): We used shRNA to specifically deplete the expression of Ncan in mPFC neurons of 2-to 3-month-old mice after confirming knockdown efficiency in cortical cultures. We then studied mPFC-dependent forms of learning and memory in Ncan shRNA-injected mice and observed a loss of temporal order recognition memory and impairment in reversal spatial learning in these animals. Finally, using immunohistochemistry and RT-qPCR to unravel potential cellular mechanisms, we observed a significant reduction in the perisomatic GABAergic innervation of PV+ cells in both Ncan KO and Ncan shRNA-injected mice.

## METHODS

### Animal Housing and Ethics

All animal experiments were conducted in accordance with the ethical animal research standards defined by German and Spanish law with the Directive 2010/63/EU of the European Parliament and of the Council of 22 September 2010 on the protection of animals used for scientific purposes and approved by the Ethical Committee on Animal Health and Care of the State of Saxony-Anhalt, Germany, with license numbers 42502-2-1322 DZNE and 42502-2-1343 DZNE, or by the Committee on Bioethics of the Universitat de València. This study used a total of 30 male mice including five wild-type and five *Ncan* KO 3-month-old male mice derived from the same founder colonies (Zhou et al., 2001), ten 2- to 3-month-old male C57BL6/J mice injected with Control shRNA AAV (hereafter referred to as Control shRNA), and ten 2- to 3-month-old male C57BL6/J mice injected with Ncan shRNA AAV (hereafter referred to as Ncan shRNA). The number of mice used in each experiment is given in the text and figure legends. Mice used for AAV injections were transferred to the research facility from the animal breeding house and housed individually with food and water available *ad libitum* for at least 72 hours before experiments under a reversed 12/12 light/dark cycle (light on 9 P.M.). All behavioral experiments were performed during the dark phase of the cycle, i.e. when mice are active, under constant temperature and humidity. All wild-type and *Ncan* KO mice were housed in groups of 2 to 4 in a standard environment (12 h light/dark cycle) and with *ad libitum* access to food and water. Every effort was made to minimize the number of animals used and their suffering.

### Knockdown experiments using Ncan shRNA

To knock down mouse Ncan (GeneID: NM_007789.3), shRNA plasmids were cloned by the insertion of the siRNA sequences (Sigma) shown to be effective against Ncan mRNA (Okuda et al., 2014) (Suppl. Table 1) targeting the open reading frame into AAV U6 GFP (Cell Biolabs Inc., San Diego, CA, USA) using BamH1 (New England Biolabs, Frankfurt a.M., Germany) and EcoR1 (New England Biolabs, Frankfurt a.M., Germany) restriction sites. shRNAs were designed for targeting Ncan messenger RNA (mRNA) (Ncan shRNA) by selecting target sequences in the coding regions of Ncan to prevent or minimize off-target effects (Taxman et al., 2006). The shRNA universal negative control (Sigma, Control shRNA) was used as a non-targeting control (Suppl. Table 1).

Positive clones were sequenced and used for the production of recombinant adeno-associated particles as described previously (Mitlohner et al., 2020). HEK 293T cells were transfected using PEI (1ng/ul) with an equimolar mixture of the shRNA-encoding AAV U6 GFP, pHelper (Cell Biolabs Inc., San Diego, CA, USA) and RapCap DJ plasmids (Cell Biolabs Inc., San Diego, CA, USA). From 48-72 hours after transfection, freeze-thaw cycles were implemented to lyse cells and then with benzonase (50 U/mL; Merck Millipore, Burlington, MA, USA) for 1 h at 37 °C. Lysates were centrifuged at 8000× g at 4 °C, supernatants collected, and filtered with a 0.2-micron filter. We then purified filtrates using the pre-equilibrated HiTrap Heparin HP affinity columns (GE HealthCare, Chicago, IL, USA), followed by washing with the following buffers in sequence; Wash Buffer 1 (20 mM Tris, 100 mM NaCl, pH 8.0; sterile filtered) and wash buffer 2 (20 mM Tris, 250 mM NaCl, pH 8.0; sterile filtered). Elution buffer (20 mM Tris, 500 mM NaCl, pH 8.0; sterile filtered) was then used to elute viral particles. Finally, Amicon Ultra-4 centrifugal filters with 100,000 Da molecular weight cutoff (Merck Millipore, Burlington, MA, USA) along with 0.22 µM Nalgene® syringe filter units (sterile, PSE, Sigma-Aldrich, St. Louis, MO, USA) were used to further purify viral particles before being aliquoted and stored at -80 °C.

### Cell culture, neuronal infection, RT-qPCR and GI-LTP

Cortical neurons were isolated from embryonic C57BL6/J mice (E18), as described previously (Seibenhener and Wooten, 2012). After harvesting, neurons were plated on poly-l-lysine-coated (Sigma; molecular weight, 30 kDa) in 6 well plates without coverslips at a cell density of 500,000 cortical neurons per well. Neurons were maintained in 2.5ml of Neurobasal media (Invitrogen) supplemented with 2% B27 and 1% L-glutamine and 1% Pen-Strep (Life Technologies). Dissociated cortical neurons were infected with Control and Ncan shRNA AAVs at 7 days *in vitro* (DIV) and mRNA expression levels of Ncan were assayed at DIV-21. Knockdown efficiency of Ncan shRNA normalized to Control shRNA was determined using RT-qPCR. From cortical cultures, total RNA was extracted using the EURx GeneMatrix DNA/RNA Extracol kit (Roboklon, Cat. No. E3750) according to the manufacturer’s recommendations (Ventura Ferreira et al., 2018; Baidoe-Ansah et al., 2019) and products were further checked for genomic DNA contamination by using Nano-drop and gel electrophoresis to measure the RNA yield, purity, and integrity, respectively. Then, 2 μg of RNA was used for cDNA conversion using the High-Capacity cDNA Reverse Transcription Kit (Cat. No. 4368814). qPCR analysis of 12 genes was performed with TaqMan™ gene expression array assay (ThermoFisher Scientific, Cat. No. 4331182) using the Quant-Studio-5 (Applied Biosystems). Details of all TaqMan™ probes used are given in Suppl. Table 2. Finally, gene expression per sample was normalized relative to the expression of glyceraldehyde 3-phosphate dehydrogenase (GAPDH) (Meldgaard et al., 2006). For the glycine-induced form of chemical LTP (GI-LTP), already shRNA AAV infected cortical cultures were first incubated with sterile artificial cerebrospinal fluid (ACSF) containing the following (in mM): 119.0 NaCl, 1.3 MgSO_4_ 1.0 NaH_2_PO_4_, 26.2 NaHCO_3_, 2.5 KCl, 2.5 CaCl_2_, 11.0 glucose, 0.2 glycine, for 5 minutes at 37°C before returning to ACSF without glycine for 50 minutes (Fortin et al., 2010), whereas control cultures were incubated in ACSF.

### Stereotaxic injection of Ncan shRNA and Control shRNA into the PFC

Twenty 2- to 3-month-old mice were briefly sedated with isoflurane in a chamber and fixed to a stereotactic frame (SR-6M, Narishige Scientific Instrument Lab, Japan). Mice were anesthetized with isoflurane adjusted to 4% for induction and then reduced to 1.5-2%, with oxygen levels set to 0.4 L/min (Baxter 250ml Ch.-B.: 17L13A31). The body temperature of mice was maintained at 37°C using a heating pad (ATC1000 from World Precision Instrument, USA). Ophthalmic ointment (BRAND) was applied to protect the eyes during surgery after which the skin was cleaned with 70% ethanol and hair shaved. A volume of 1000 nl was injected using 10 μl NanoFil syringe (World Precision Instrument, USA) and a calibrated glass microelectrodes connected to an Ultra microinfusion pump (UMP3, World Precision Instrument, USA) at a rate of 3 nl/sec. The coordinates for injection were dorso-ventral (DV) from the brain surface, anterior-posterior (AP) from bregma, and medio-lateral (ML) from the midline (in mm): AP, +1.77; ML, -0.3; DV, -2.2. Ncan shRNA (3.76×10^11^) and Control shRNA (5.86×10^11^) were injected bilaterally into the mPFC. After all procedures, the animals were placed in a recovery chamber under red light for 15 minutes.

### Open field and recency task

Before behavioral testing, mice were handled and habituated to the behavioral room using the glass tunnel method of handling for 3-4 days (Gouveia and Hurst, 2017). On the 5^th^ day, the open-field task was performed, where mice were placed in an empty arena (50 x 50 x 30 cm) for 10 minutes (Holter et al., 2015; Kaushik et al., 2018; Baidoe-Ansah et al., 2019). The position of animals was tracked using an overhead camera and the tracking software (Anymaze 4.99: Stoelting Co., Wood Dale, IL, USA). The total distance traveled and times spent in the central vs. peripheral areas were used as indicators of mice’s general activity and anxiety. The temporal aspect of learning and memory has been shown to depend on mPFC, a region of the CNS associated with executive function. One of the behavioral tasks capable of assessing this aspect is the temporal order recognition (recency) task (Naya et al., 2017). It was performed 24 hours after the open-field task. Before and after each session, the floors and walls were cleaned with 70% ethanol. In each of three sessions, mice were placed at the center of the apparatus using the glass tunnel – initially facing away from two objects – and allowed to explore for 10 minutes. During the first session, two indistinguishable objects were presented to the mice (First-Sampling-phase: objects **A, A**). The second set of different objects (Second-Sampling-phase: **B, B**) was presented after an hour. Fifteen minutes later, a probe test was performed with one object from each training session being presented (**A**, **B**). The time intervals spent exploring these objects (less-recently-experienced; **A** and recently-experienced; **B**) were measured by a trained observer who was blinded to treatment conditions (Bonardi et al., 2016). Exploring time for old (less-recently-experienced) and recent (recently-experienced) objects, as well as discrimination ratio [(old-recent)/ (old+recent)] x 100 %, was used to evaluate animals’ recognition memory.

### Labyrinth task (Dry maze)

The Labyrinth task also known as the dry maze is a behavioral setup capable of assessing hippocampus-dependent spatial learning and memory, which can also be used to test mPFC-dependent memory retrieval and reversal learning (Peters et al., 2013). This setup is non-stressful, compared to the water maze (Harrison and Feldman, 2009), yet capable of measuring the various learning strategies mice use for spatial navigation, such as place and response learning, and the egocentric strategy (Rich and Shapiro, 2007; Peters et al., 2013). Using the labyrinth set-up to study mPFC functions, we designed a 3-phase-learning paradigm that included an initial-learning-phase (4 days), 2 relearning-phases (3 days each) and a 3-day inter-learning delay (Figure 3A&B).

Before the labyrinth task, mice were restricted to 1.0 ml of water per day for a maximum of 5 days. On the 6^th^ day (day 1 of the initial-learning-phase), mice were allowed to explore the maze for 10 min with 1.0 ml of water (as a re-enforcer) located in the reward zone. Mice were then placed in the maze for 3 trails per day on days 7, 8, 13, 14, 20 and 21; with each trial manually ended upon entry into the reward zone. An inter-trial delay of 1 hour was used for each training session. On days 9, 15, and 22, a probe test was performed, which included a 90° rotation and dropped curtains to enclose the maze and exclude extra-maze (distal) cues. This particular probe test was introduced to test the use of the allocentric vs egocentric strategies of navigation. During the relearning-phase, the reward path (Figure 3B, green line) and reward location (green zone) were changed while maintaining the start zone. A decision point was generated by blocking the recently learned path toward the reward. A learning curve was then measured using an overhead camera with animal tracking software (Anymaze 4.99: Stoelting Co., Wood Dale, IL, USA). The following parameters; distance traveled before entry into the reward zone, the number of errors (indicated by red path line), and latency to reward, were calculated. For all phases of learning, the start zone was kept constant. During the relearning phases, mice were exposed to a redesigned-labyrinth (new path and reward location). This was to measure the mPFC function in decision making as shown by Peters and colleagues (Peters et al., 2013). The old path was blocked (gray zones; **Zone-1** for 1^st^ re-learning and **Zone-1&2** for 2^nd^ re-learning) and the contribution of mPFC to decision-making was assessed at the **decision point**; i.e. the point of intersection between old and new paths (Figure 3A&B).

Moreover, several studies have used within-day and between-day error indices to show neural dissociations for spatial navigation (Churchwell et al., 2010). Therefore, in this study, we used the mean number of errors to measure the between-day retrieval index, which would reflect overnight consolidation of acquired memories, as the percentage difference in performance between the two training sessions with a 22 ± 2 hour-interval between them, i.e. between t3 (Er*_t3_*) on day 7, 13, or 20 and t1 (Er*_t1_*) on day 8, 14, or 21 (Vago et al., 2007). Next, to measure the decision accuracy index, we determined the difference between the number of entries into the blocked zones (gray zones, Figure 3B) for both reversal learning phases. Here, the encoding score (En*_e_*), as illustrated below, was operationally defined in this study as the mean number of entries into a blocked path across the last two training sessions on a particular day (i.e. t2 and t3 on days 13 and 20), whereas the first training session 22 ± 2 hours later was selected as retrieval score (E*_r_*) (Churchwell et al., 2010). The percentage difference (% Accuracy index) was then calculated. The following formulas were then used:

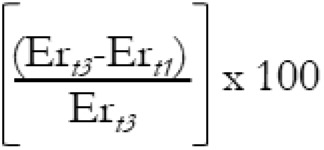

1. 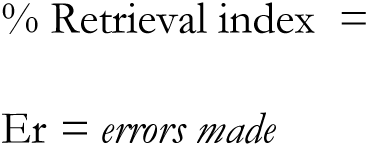
2. 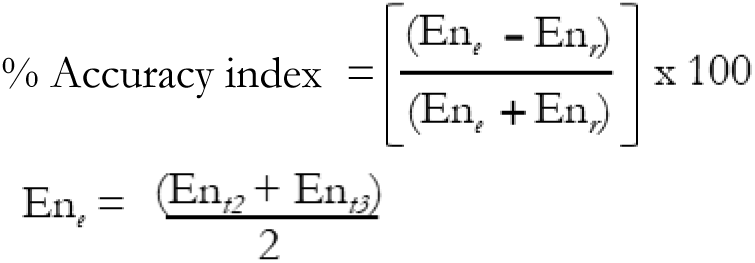

En = *entry into the blocked path*

En*_e_* = *encoding scores for entry into the blocked path*

En*_r_* = *retrieval scores for entry into the blocked path*

### Immunohistochemistry

Ncan and Control shRNA-injected mice were anesthetized using isoflurane and perfused transcardially with ice-cold phosphate-buffered saline (PBS) and fixed with 4% PFA for 10 min. Mice brains were then dissected and incubated in 4% PFA overnight and cryoprotected in 30% sucrose for 48 h, and frozen in 100% 2-methylbutane at -80°C. The dissected brains were incubated in 4% PFA-containing PBS, cryoprotected in 30% sucrose-containing phosphate buffer (PB) solution for 48 hours, and frozen in 100% 2-methylbutane at -80°C. Slices of 50-μm thick coronal sections were kept floating in solution (1-part ethylene glycol, 1-part glycerin, 2 parts PBS, pH = 7.2) at 4°C. For each staining, at least 2 sections per animal were used. The sections were first washed in 120 mM phosphate buffer (PB) at a pH of 7.2 and then permeabilized with PB containing 0.5% Triton X-100 (Sigma, T9284) for 10 min at 37 °C. Then sections were blocked with a blocking solution (PB supplemented with 0.3% Triton X-100 and 5% normal goat serum, NGS, Gibco, 16210-064) for 1 h at 37°C. Sections were then incubated with primary (for 20 h) and secondary antibodies (for 3 hr) at 37°C and were washed 3 times in PB after each incubation. Finally, sections were mounted on Superfrost glasses with Fluoromount (Sigma, F4680) and then confocal images were taken with the Zeiss LSM 700 microscope.

The wild-type and Ncan KO mice were perfused as follows. Deep anesthesia was induced by isoflurane overdose, then animals were perfused transcardially, first with NaCl 0.9% for 1 minute and then with PFA 4% in phosphate buffer (PB) 0.1 M for 20 min. After perfusion, brains were extracted and post-fixed by immersion in PFA 4% for 2 h. Then, brains were cryoprotected in 30% sucrose in PB 0.1 M and frozen. Afterward, brains were cut into 40 microns’ sections using a cryostat, and sections were collected and stored at -20°C. Sections were processed for fluorescence immunohistochemistry as follows. Slices were washed in saline phosphate buffer (PBS), then they were incubated for 2 h in 5% normal donkey serum (NDS, Gibco), 0.2% Triton-X100 in PBS to block nonspecific binding. After that, sections were incubated for 48 h at 4°C with different primary antibodies (Suppl. Table 2). Then, sections were washed and incubated for 2 hours at room temperature with secondary antibodies (Suppl. Table 3). Finally, sections were washed in PB 0.1 M, mounted, and cover-slipped using Vectashield fluorescence mounting medium (Vector Laboratories).

### Data acquisition, processing, and analysis

Two Control shRNA mice were excluded using the outlier function (ROUT method Q = 1%) in GraphPad Prism 8.0 (GraphPad Software Inc., La Jolla, USA) after the 2^nd^ recency task whereas one Ncan shRNA mice died after 1^st^ recency task. Additionally, one Control shRNA was further excluded due to significantly low or no GFP expression. For the vGLUT1+ puncta analysis, two Ncan shRNA sections were excluded since sections and staining were of low quality. Overall, sections from 7 Control shRNA and 9 Ncan shRNA-treated mice were used for the IHC analysis. Sections from Ncan shRNA-injected and Ncan KO mice were analyzed under a confocal microscope with the Zeiss LSM 700 microscope and Leica SPE, respectively. Stacks were obtained from layer V of the PrL at an optimal penetration depth in the tissue (up to 6 μm deep). We used a 63x objective plus 2x digital zoom for the imaging of perisomatic and neuropil puncta. The acquiring conditions were maintained throughout all the imaging sessions to compare the fluorescence intensity between samples. For the quantification of perisomatic puncta in shRNA-treated mice, on PV+ and PV-cells, we outlined manually the profile of the cell somata and created a band of 1.5 μm as the region of interest (ROI). The bands were duplicated, cleared outside, and the background was subtracted with the rolling value of 50 and Gaussian blur with the s-value of 1 using the find maxima plugin in Fiji with a prominence of 5, size (0.3 to 1.0 μm^2^) and circularity (0.5 – 1.0), we measured the mean intensity along with the number of synaptic puncta. For neuropil analysis, we randomly selected 4 rectangular ROIs (100μm^2^) per image and used the aforementioned procedure to measure the fluorescent intensity and the number of synaptic puncta. Additionally, analysis of images from Ncan KO mice was performed as previously described (Schindelin et al., 2012; Guirado et al., 2018). Briefly, after applying the subtract background tool (the rolling value = 50) and Gaussian blur (s value = 1) filters, the original outline was expanded 1 μm from the cell body edge and the region of interest (ROI) was defined as the area between both outlines. Then, ROI was segmented and binarized using a dynamic threshold. The number of puncta was quantified after filtering particles for size (included from 0.3 to 1 μm^2^) and circularity (included from 0.5 to 1). Finally, the selection of binarized puncta was overlapped on the original image to measure the fluorescence intensity (FI) of puncta. For the analysis of neuropil puncta, ten squares (100 μm^2^) were randomly selected for analysis. Images were processed using the previously explained methodology.

### Statistical analysis

Statistical analysis was performed using GraphPad Prism 8.0 (GraphPad Software Inc., La Jolla, USA) and Statistica 8.0 (StatSoft, USA) software. All data are shown as mean ± SEM with n being the number of mice. Asterisks in figures indicate statistical significance (with details in the figure legend or Results). The hypothesis that experimental distributions follow the Gaussian law was verified using Kolmogorov-Smirnov, Shapiro-Wilk, or D’Agostinio tests. For pairwise comparisons, we performed the Students’ t-test where the samples qualify for the normality test; otherwise, the Mann-Whitney test was employed. Additionally, Wilcoxon matched-pairs test was used for paired data that did not pass the normality test. Holm-Sidak’s multiple comparisons t-test was used for independent comparisons. Spearman correlation coefficients were computed to estimate the correlation between the variables studied. As indicated, one- and two-way ANOVA with uncorrected Fisher’s LSD as well as Brown-Forsythe with Welch ANOVA tests were also used. The p-values represent the level of significance as indicated in figures by asterisks (*p < 0.05, **p < 0.01, ***p < 0.001 and **** p < 0.0001) unless stated otherwise.

## RESULTS

### Validation of Ncan shRNA

To study the contribution of Ncan to the maintenance of PNNs around PV+ interneurons as well as in cognitive and synaptic functions, we evaluated AAV encoding shRNA targeting the Ncan mRNA. The knockdown efficiency was estimated by using RT-qPCR and immunohistochemistry (Figure 1A, B, D, E & F). We infected cortical cultures with the Ncan shRNA and Control shRNA vectors at DIV7, and determine Ncan mRNA expression levels at DIV21 using qRT-PCR (after normalization to the GAPDH level as an internal control) (Meldgaard et al., 2006; Baidoe-Ansah et al., 2019). We observed that treatment with Ncan shRNA resulted in about 80% reduction (p < 0.0001, Figure 1A).

**Figure 1.**
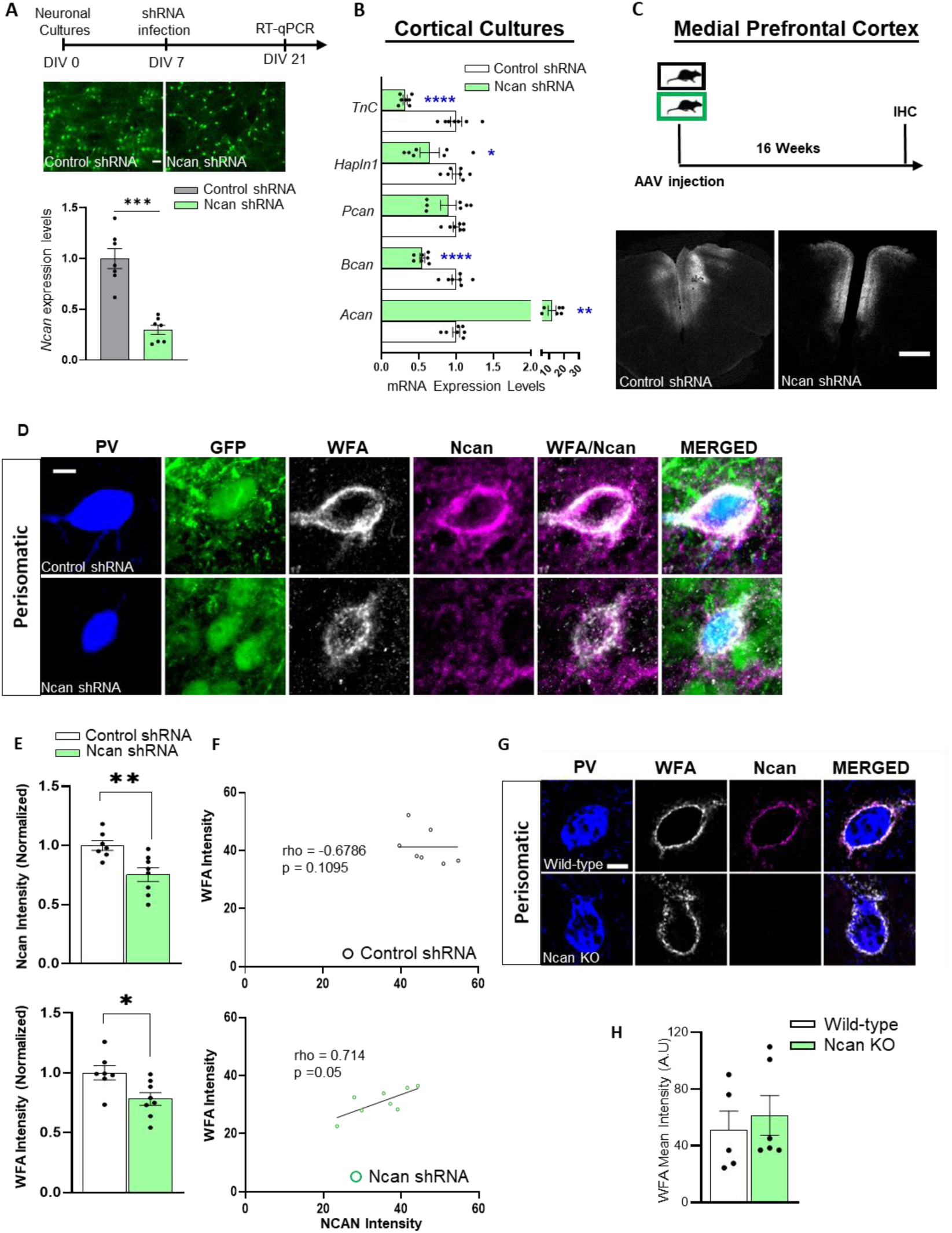
Knockdown of Ncan expression in the PFC affected PNN maintenance. The shRNA constructs developed to knockdown Ncan were characterized in cultured cortical neurons, and in the PFC sections after injection of shRNA AAVs in 2- to 3-month-old mice. **(A)** Timeline for *in vitro* validation of Ncan knockdown in cultured cortical cells by RT-qPCR. The representative images of Ncan and Control shRNA infected cortical cells expressing GFP. Scale bar 100 μm. **(B)** Ncan knockdown upregulated Acan and downregulated Bcan, HAPLN1 and TnC in cultured cortical neurons. **(C)** A scheme of experiments and examples of AAV expression 16 weeks after injection in the infralimbic part of PFC. Scale bar 100 μm. **(D)** Representative images of PNNs that were visualized and quantified after Ncan knockdown. Scale bar, 5 μm. **(E)** The perisomatic expression of WFA and Ncan per animal was reduced in Ncan shRNA AAV-injected mice compared to Control shRNA AAV-injected mice. **(F)** Additionally, a positive correlation between the expression of Ncan and WFA in Ncan shRNA-treated mice but not in controls. Three-month-old Ncan KO mice were compared to aged-matched wild-type mice. **(G)** Using WFA and Ncan antibodies, we show a lack of Ncan in PNNs of Ncan KO mice in representative images. Scale bar, 5 μm. **(H)** Quantification of the mean immunofluorescence intensity of WFA in Ncan KO mice. Bar graphs show mean ± SEM values. *p < 0.05, **p < 0.01, ***p < 0.001 and **** p < 0.0001, represent significant differences between Control shRNA (N = 7) and Ncan shRNA (N = 9) treated mice or wild-type (N = 5) and Ncan KO (N = 5) mice using one-way ANOVA with Sidak’s multiple comparisons test; Brown-Forsythe and Welch ANOVA tests; unpaired t-test with Welch’s correction, or significant Spearman coefficients of correlation.

Next, we used shRNA-infected cortical cultures to investigate the effect of Ncan reduction on the expression of other ECM molecules and found HAPLN1 and Bcan mRNA levels to be downregulated (p = 0.034 and p < 0.0001, respectively; Brown-Forsythe and Welch ANOVA tests, Figure 1B), in agreement with a recent study of Ncan KO mice (Schmidt et al., 2020). Additionally, we found a downregulation of the Ncan-binding glycoprotein Tenascin-C (TnC) (Grumet et al., 1994; Zhou et al., 2001) (p < 0.0001 Brown-Forsythe and Welch ANOVA tests, Figure 1B), whereas no change in expression of phosphacan/PTPRz1 (Pcan) occurred. Strikingly, Acan mRNA was strongly upregulated in Ncan shRNA-infected cells (p = 0.0059; Brown-Forsythe and Welch ANOVA tests, Figure 1B). These findings support the view that Ncan may be essential for the overall expression of multiple ECM molecules.

### Knockdown of Ncan expression impaired PNN maintenance

We then injected Ncan shRNA AAV into infralimbic PFC (corresponding to mPFC in human) of 2-3-month-old C57BL6J male mice to study the *in vivo* effects of Ncan depletion (Figure 1C). Using immunohistochemistry, we compared the perisomatic expression of both Ncan and WFA labeling of PNNs around PV+ interneurons in mPFC after either knockdown or knockout of Ncan (Figure 1D). We analyzed the fluorescent intensity of WFA and Ncan expression per PV+ cell and observed no significant difference for WFA but a reduction for Ncan (p = 0.377 and p = 0.008, respectively; nested t-test, Figure S1B). Analysis of mean values per animal confirmed that the expression of Ncan shRNA significantly reduced Ncan protein expression (p = 0.0046; unpaired t-test with Welch’s correction, Figure 1E). This was associated with a significant decrease in the expression of WFA (p = 0.049; Unpaired t-test with Welch’s correction, Figure 1E). Moreover, we observed a significant positive relationship between the expression of Ncan and WFA in the Ncan shRNA groups of animals and cells (Spearman coefficient of correlation rho = 0.714; p = 0.05 and rho = 0.559; p < 0.0001, respectively; Figure 1F & S1C), but no such relationship occurred for Control shRNA, suggesting that the forced downregulation of Ncan impaired the maintenance of the molecular composition of PNNs. Then, we examined the effect of constitutive Ncan KO on WFA expression in PNNs (Figure 1G) and surprisingly (but in agreement with Zhou et al., 2001) observed no significant difference between groups (p = 0.608; unpaired t-test with Welch’s correction, Figure 1H). In these experiments, we also confirmed the specificity of the Ncan antibody by showing a lack of expression of Ncan in the PFC of Ncan KO mice (Figure 1G).

### Impaired temporal order recognition and reversal spatial learning after Ncan knockdown

Next, we aimed to determine the cognitive effects of Ncan knockdown in the PFC. Therefore, we measured two essential PFC-dependent cognitive functions, namely temporal order recognition memory and spatial reversal learning, by using the recency test and the labyrinth (dry maze) task, respectively. Locomotor behavior and anxiety were controlled by subjecting mice to the open field test. Considering the slow turnover of mature ECM components (Decaris et al., 2014; Matsubayashi et al., 2020), we performed tests 8 and 16 weeks after Ncan shRNA AAV delivery (Figure 2A). In the open field test, there was no difference in the distance traveled between these treatments neither 8 nor 16 weeks after AAV injection (p = 0.492 and p = 0.569, respectively; Unpaired t-test with Welch’s correction, Figure 2B_2_ & 2C_2_). In the temporal order recognition task we observed no significant difference between Ncan shRNA and Control shRNA animals after 8 weeks (discrimination index: 24.43 ± 5.18 % vs 15.93 ± 6.07%, respectively; p = 0.315; Mann-Whitney U-test, Figure 2B_3_). However, 16 weeks after Control shRNA AAV injection, mice explored significantly longer the less recently seen (old) objects than the most recently (recent) presented objects with exploration times 55.24 ± 7.871 s vs. 19.08 ± 1.349 s, respectively (p = 0.007; Wilcoxon matched-pairs test, Figure 2C_3_). In contrast, mice injected with Ncan shRNA AAV failed to discriminate between both objects (39.34 ± 3.520 s vs. 36.17 ± 6.933 s; p = 0.496; Wilcoxon matched-pairs test, Figure 2C_3_). A discrimination ratio analysis further confirmed the difference between Control shRNA and Ncan shRNA groups (44.87 ± 6.624 % vs 9.380 ± 11.75 %; p = 0.027; Mann-Whitney U test, Figure 2C_3_). These results suggest that the loss of Ncan in the PFC impairs temporal order recognition memory.

**Figure 2.**
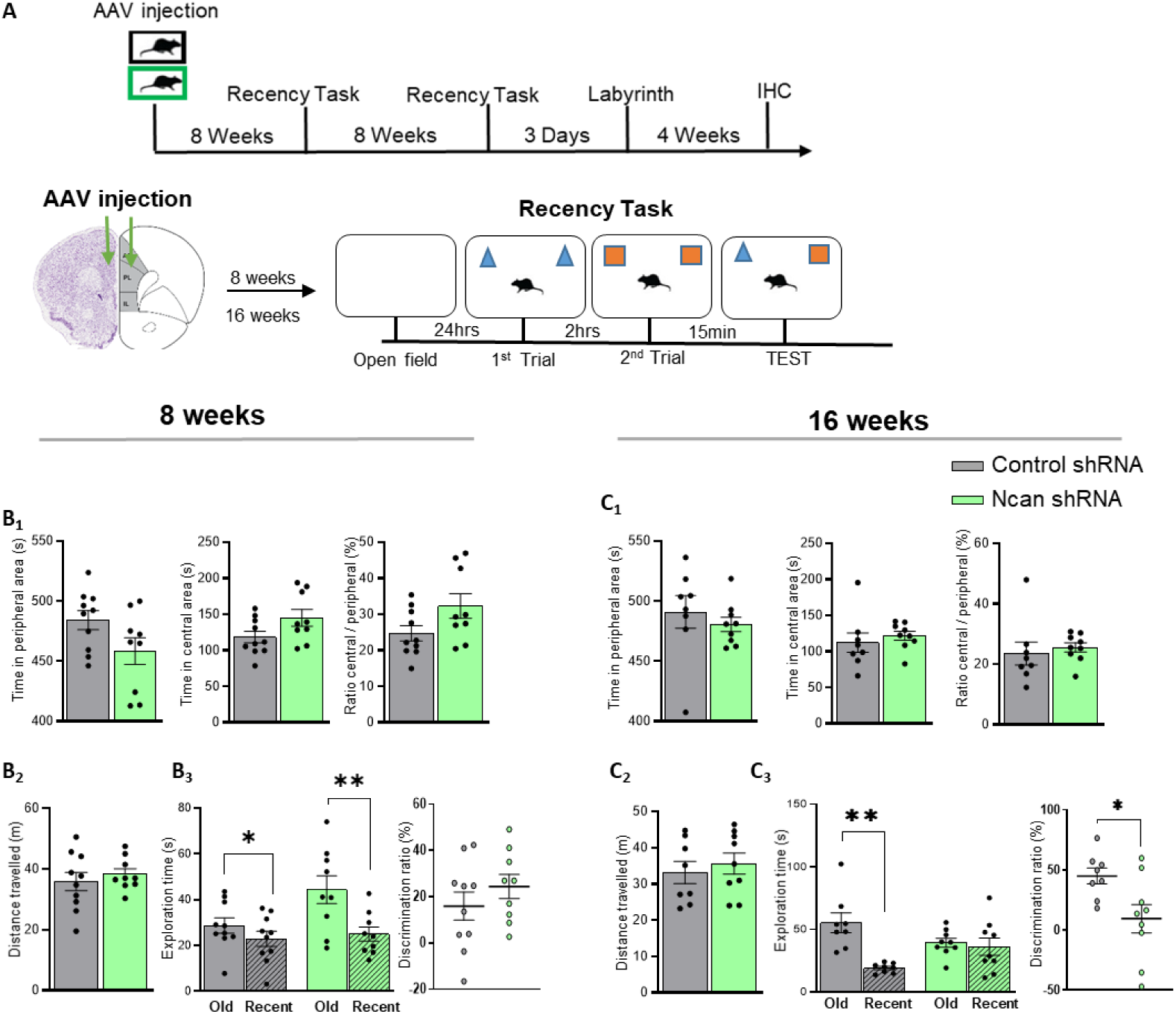
Knockdown of Ncan expression in the PFC impaired temporal order recognition memory. The PFC-dependent recency task was performed in mice injected with Ncan shRNA and Control shRNA AAVs. **(A)** Timeline of behavioral experiments and the recency task used in this study. The time intervals spent exploring old and recent objects 8 weeks **(B)** or 16 weeks **(C)** after injection of AAV were measured, with the significant effect of Ncan knockdown observed after 16 weeks. Bar graphs show mean ± SEM values. *p < 0.05, **p < 0.01, ***p < 0.001 and **** p < 0.0001 represent significant differences between Control shRNA (N = 10) and Ncan shRNA (N = 10) treated mice for 8 weeks and Control shRNA (N = 8) and Ncan shRNA (N = 9) for 16 weeks using Mann-Whitney U-test; unpaired t-test with Welch’s correction and Wilcoxon matched-pairs test.

Then, we investigated the effects of Ncan knockdown on spatial learning and reversal learning (relearning) in the Labyrinth considering that the decision-making during relearning (cognitive flexibility) critically depends on the mPFC (Guise and Shapiro, 2017) (Figure 3A&B). Overall, no difference in distance traveled (p = 0.735; two-way RM ANOVA, Figure 3C) and mean number of errors made (p = 0.724; two-way RM ANOVA, Figure 3D) was observed during the learning and relearning phases. However, Ncan shRNA-treated mice traveled longer distances and made significantly more errors at the 2^nd^ training session on day 13 during the 1^st^ relearning phase (p = 0.0406 and p = 0.0286, respectively; two-way ANOVA with Fisher’s LSD test, Figure 3C&D). This observation suggests a potential decrease in short-term encoding and retrieval efficiencies due to Ncan shRNA, although this effect was lost in the 2^nd^ relearning phase. We also estimated the retrieval index (as defined in methods) between the last training session (t3) on the first day and the first session (t1) of the next day for each phase, by using the mean number of errors. During the learning phase, we observed no difference in the retrieval of spatial memory between t3 on day 7 and t1 on day 8 (47.50 ± 14.00 % vs 41.99 ± 21.69 %; p = 0.8344; unpaired t-test with Welch’s correction, Figure 3D) for both groups of animals. There was a weak tendency of impaired retrieval of long-term reversal spatial memory in Ncan shRNA i.e. during the 1^st^ relearning phase (14.79 ± 8.5570 % vs -3.831 ± 10.06 %; p = 0.1282; unpaired t-test with Welch’s correction, Figure 3D). However, the retrieval performance of Ncan shRNA-treated mice showed a strong tendency to be impaired as compared to controls during the 2^nd^ relearning phase (32.05 ± 8.196 % vs -1.358 ± 14.09 %; p = 0.0617; unpaired t-test with Welch’s correction, Figure 3D). Additionally, we used the accuracy index (as defined in methods) to measure the accuracy of the first decision-making where to go, and we found mice injected with Ncan shRNA to make more errors at the decision point by entering the blocked paths in both relearning phases. This occurred 22 ± 2 hours after mice relearned the new path to the new reward location, i.e. exactly at t1 on both day 14 (1^st^ relearning-phase; 37.50 ± 18.30 % vs -7.407 ± 7.407 %; p = 0.0482; unpaired t-test with Welch’s correction, Figure 3E) and day 21 (2^nd^ relearning-phase; 46.40 ± 11.20 % vs 13.60 ± 12.32 %; p = 0.0676; unpaired t-test with Welch’s correction, Figure 3E). This result suggests that Ncan may be essential for the accurate retrieval of reversal spatial memory after its overnight consolidation.

**Figure 3.**
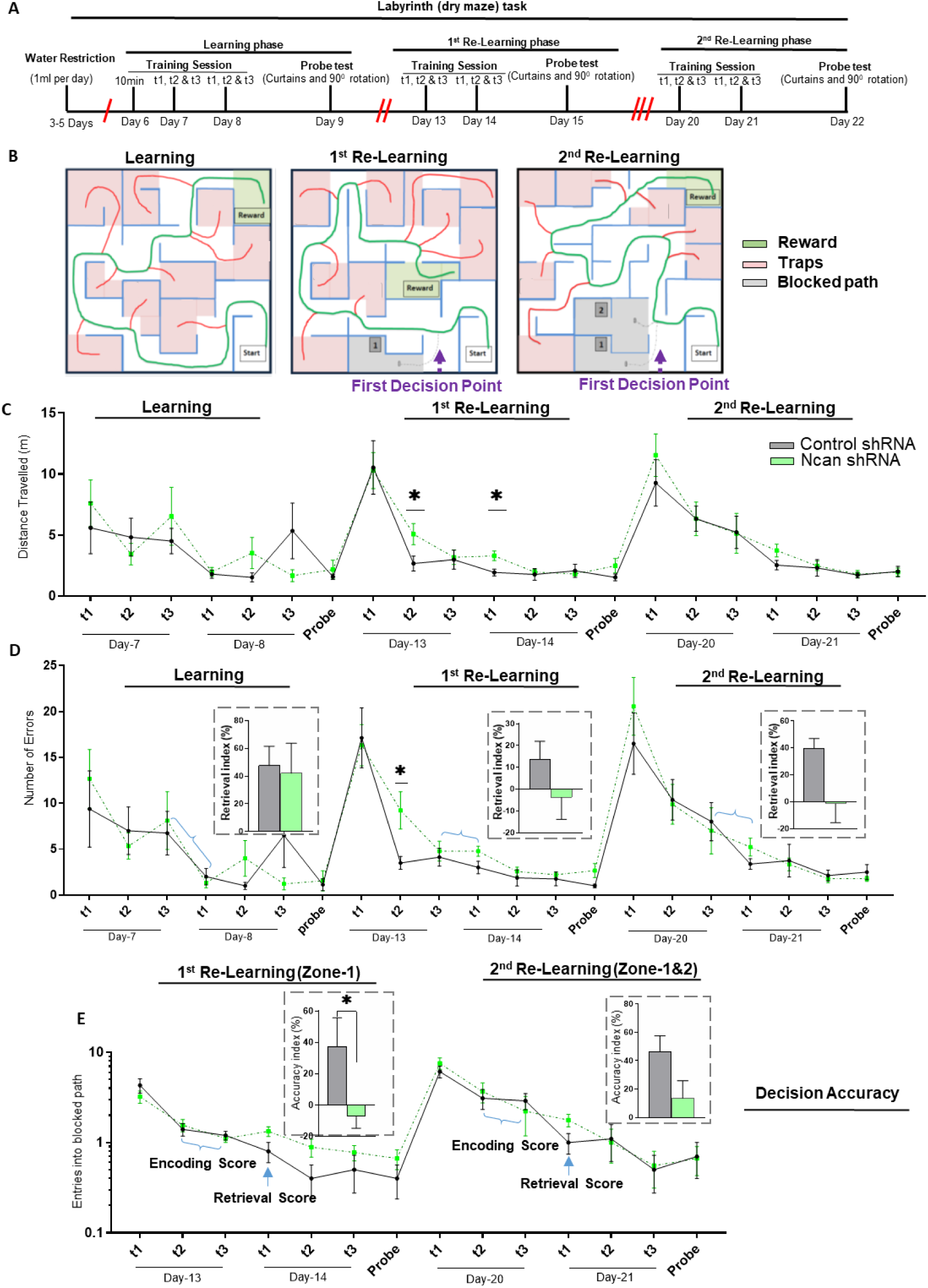
Ncan knockdown impaired consolidation of spatial memory after reversal learning. The Labyrinth (dry maze) task was implemented using water as a reward. **(A)** Timeline for the labyrinth task that was designed to include one phase of learning and two phases of reversal learning. **(B)** Design of the labyrinths used, pointing to the start zone, a reward location (green zone), traps (pink zones), blocked paths (gray zones) along the right paths (green line) and wrong paths (red lines). The performance of mice was measured using the distance travelled and the number of errors made. **(C)** No difference in the mean distance travelled between Control and Ncan shRNA-treated mice was observed. **(D)** Ncan shRNA-treated mice potentially failed to consolidate spatial memory during the two re-learning phases of the Labyrinth task. **(E)** Ncan shRNA-treated mice were less accurate in discriminating between the blocked path (s) and the new path to the new reward location. Bar graphs show mean ± SEM values. *p < 0.05, **p < 0.01, ***p < 0.001 and **** p < 0.0001 represent significant differences between Control shRNA (N = 8) and Ncan shRNA (N = 9) treated mice using two-way RM ANOVA; two-way ANOVA with Fisher’s LSD and unpaired t-test with Welch’s correction.

### Reduced GABAergic innervation after Ncan knockdown

Prompted by the effect of Ncan dysregulation on temporal recognition memory and reversal spatial learning, as observed in this study, we then analyzed the possible cellular and molecular underpinnings. We used immunohistochemistry to quantify synaptic changes around PFC inhibitory and excitatory neurons in Ncan shRNA-treated and Control shRNA-treated mice, as previous studies have suggested a potential role of Ncan in perisomatic innervation (Sullivan et al., 2018; Schmidt et al., 2020). Using antibodies against vesicular GABA transporter (vGAT) (Figure 4A), we found a significant reduction in the fluorescent intensity (p = 0.0137; unpaired t-test with Welch’s correction, Figure 4B) and a strong tendency of reduction regarding the density of vGAT+ perisomatic puncta (p = 0.0609; unpaired t-test with Welch’s correction, Figure 4B) on PV+ neurons of Ncan shRNA-treated mice whereas no such alterations occurred on PV-neurons for both groups. In the neuropil area (Figure 4C), Ncan depletion did not affect the intensity and the number of vGAT+ puncta (p = 0.2062 and p = 0.8930, respectively; unpaired t-test with Welch’s correction, Figure 4D). Additionally, we found a reduced fluorescent intensity (p = 0.0030 and p = 0.0052, respectively; Brown-Forsythe and Welch ANOVA tests, Figure 4B) and a reduced number of PV+ puncta (p = 0.0004 and p = 0.0197, respectively; Brown-Forsythe and Welch ANOVA tests, Figure 4B) on PV- and PV+ neurons in Ncan shRNA mice and made similar findings in the neuropil area (p = 0.0055 and p = 0.0047, respectively; unpaired t-test with Welch’s correction, Figure 4D).

**Figure 4.**
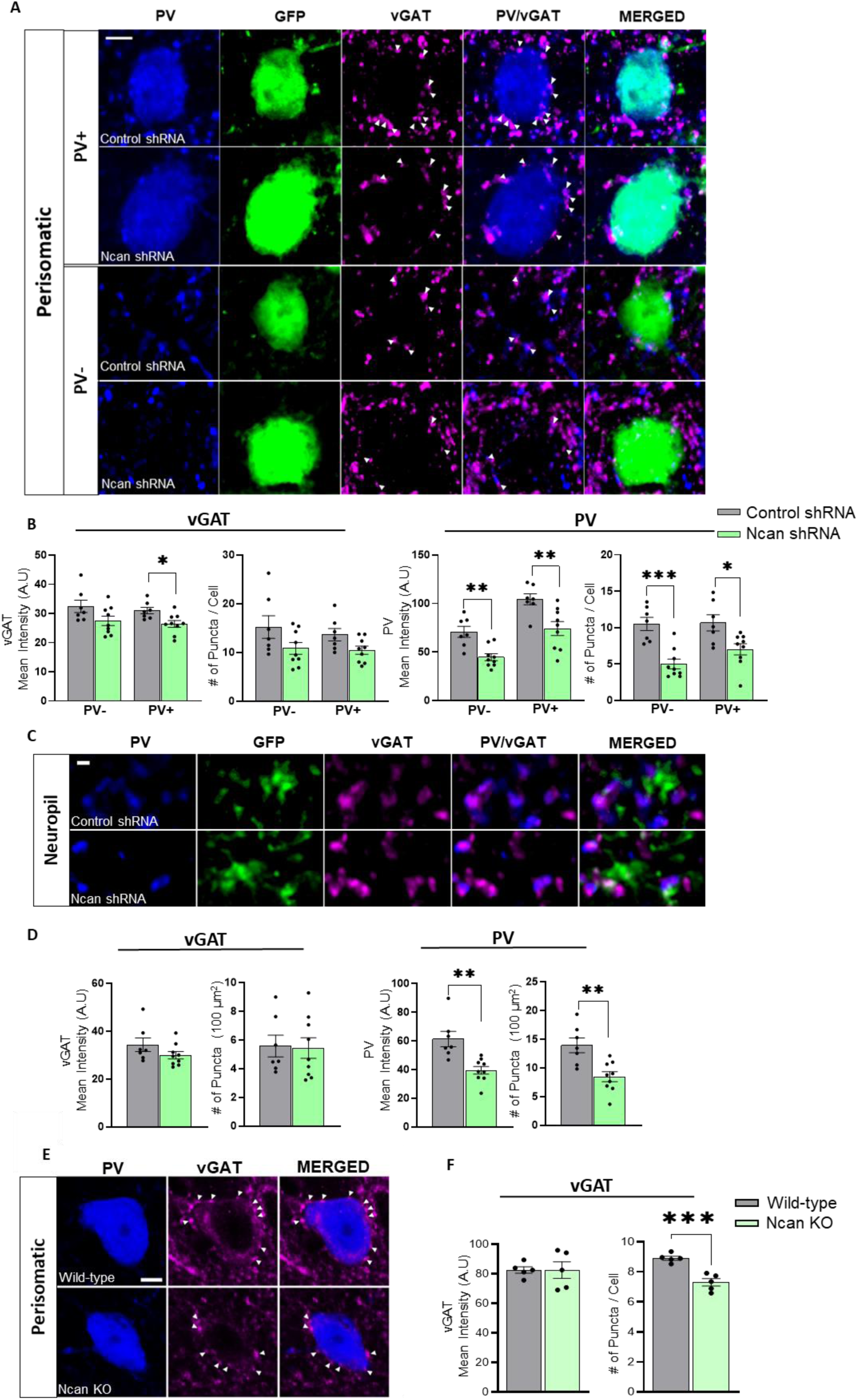
Ncan knockdown and knockout reduce perisomatic GABAergic innervation of PV+ interneurons. **(A)** Representative images showing the expression of vGAT+ puncta on shRNA-infected PV- and PV+ neurons in the PFC. White arrowheads point to vGAT+ puncta. Scale bar, 5 μm. **(B)** In Ncan shRNA-treated mice, the perisomatic fluorescence intensity of vGAT+ and PV+ puncta were significantly reduced relative to Control shRNA. **(C)** Representative images showing the neuropil expression of vGAT+ and PV+ puncta. Scale bar, 1 μm. **(D)** The fluorescent intensity and number of PV+ puncta were significantly reduced in Ncan shRNA-treated mice, but not for vGAT+ puncta. **(E)** Representative images showing vGAT+ puncta contacting the somata of PV+ interneurons in 3-month-old Ncan KO mice or aged matched wild-type mice. White arrowheads point to vGAT+ puncta. Scale bar, 5 µm. **(F)** Quantification of the mean immunofluorescence intensity and the number of vGAT puncta revealed a significant reduction of GABAergic innervation in Ncan-knockout mice. Bar graphs show mean ± SEM values. *p < 0.05, **p < 0.01, ***p < 0.001 and **** p < 0.0001 represent significant differences between Control shRNA (N=7) and Ncan shRNA (N=9) treated mice or wild-type and Ncan KO mice using Brown-Forsythe and Welch ANOVA tests and unpaired t-test with Welch’s correction.

Additionally, we quantified the perisomatic expression and number of vGAT+ and PV+ puncta on CamKII+ pyramidal neurons (Figure S2A), constituting the majority of PV-cells, and observed no difference for vGAT between groups (p = 0.3268 and 0.2145, respectively; unpaired t-test with Welch’s correction, Figure S2B). On the other hand, knockdown of Ncan decreased the fluorescent intensity and the number of PV+ puncta on CAMKII+ pyramidal neurons (p = 0.0060 and 0.0036, respectively; unpaired t-test with Welch’s correction, Figure S2C). We did not detect any difference in the fluorescent intensity of CaMKII between both shRNA groups (p = 0.2404; unpaired t-test with Welch’s correction, Figure S2D).

We then studied the perisomatic GABAergic innervation in Ncan KO mice (Figure 4E). Similar to Ncan shRNA mice, we found a reduced perisomatic number of vGAT puncta on PV+ cells of Ncan KO mice as compared to wild-type mice, whereas the fluorescence intensity of these puncta was not affected (p = 0.0006 and p = 0.9926, respectively; unpaired t-test with Welch’s correction, Figure 5F). To analyze the effects of the genetic depletion of Ncan on the perisomatic inhibition of pyramidal neurons, we quantified the number of puncta expressing vGAT and PV on CaMKII+ pyramidal neurons (Figure S2E). The number of vGAT+ puncta was normal (p = 0.5254; unpaired t-test with Welch’s correction, Figure S2F) as well as their fluorescence intensity (p = 0.9480; unpaired t-test with Welch’s correction, Figure S2F). No significant difference between wild-type and Ncan KO mice was found regarding the number of PV+ puncta (p = 0.7749; unpaired t-test with Welch’s correction, Figure S2G). Nevertheless, the fluorescence intensity of PV+ boutons surrounding CaMKII+ pyramidal neurons was significantly higher in Ncan KO mice (p = 0.0199; unpaired t-test with Welch’s correction, Figure S2G). Finally, we analyzed the fluorescence intensity of CaMKII-expressing somata and we found it significantly higher in Ncan KO mice (p < 0.0001; unpaired t-test with Welch’s correction, Figure S2H).

**Figure 5.**
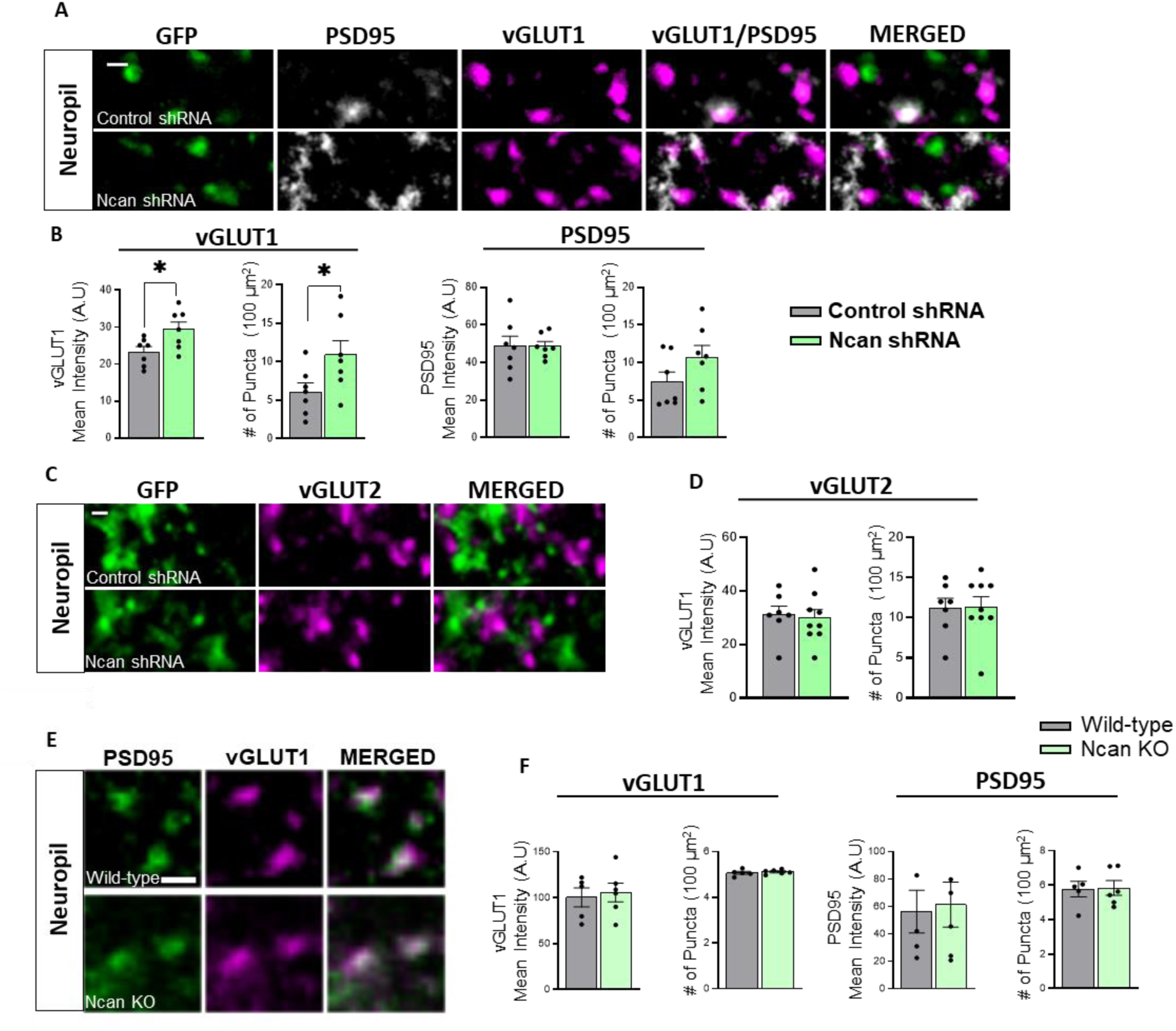
Ncan knockdown increases glutamatergic innervation by vGLUT1+ terminals. The effects of Ncan knockdown and knockout on the presynaptic expression of vGLUT1 & 2 were studied. Using antibodies against vGLUT1 and PSD95 as shown in representative images **(A)**, we quantified the density of vGLUT1+ presynaptic excitatory synapses in the neuropil area of Ncan-knockdown mice **(B).** Using antibodies against vGLUT2 as shown in representative images **(C)**, we quantified the density of vGLUT2+ presynaptic excitatory synapses in the neuropil area of Ncan-knockdown mice **(D).** The same analysis as shown in **(A, B)** for Ncan knockdown mice is shown for Ncan KO mice in **(E, F)**. Scale bar, 1 µm. Bar graphs show mean ± SEM values. *p < 0.05, **p < 0.01, ***p < 0.001 and **** p < 0.0001 represent significant differences between wild-type (N=5) and Ncan KO (N=5) mice or Control shRNA (N=7 and 7 for vGLUT1 and vGLUT2, respectively) and Ncan shRNA (N=7 and 9 for vGLUT1 and vGLUT2, respectively) using the unpaired t-test with Welch’s correction.

### Increased glutamatergic innervation after Ncan knockdown

Considering the expression of Ncan within the perisynaptic ECM of excitatory synapses, we aimed to study the effects of Ncan depletion on excitatory synapses in the PFC (Figure 5A). Hence, we analyzed the expression of presynaptic vesicular-glutamate transporter proteins (vGLUTs) in the neuropil area. The vGLUT family includes vGLUT1, vGLUT2 and vGLUT3 which are essential for the uploading of L-glutamate into synaptic vesicles of excitatory neurons (Cheng et al., 2011; Nordenankar et al., 2015). The fluorescent intensity and the number of vGLUT1+ puncta in the neuropil area were significantly higher in Ncan shRNA-compared to Control shRNA-treated mice (p = 0.0322 and p = 0.0474, respectively; unpaired t-test, Figure 5B). In contrast, the fluorescent intensity and the number of vGLUT2+ puncta (Figure 5C) were not different between shRNA groups (p = 0.959 and p = 0.957, respectively; unpaired t-test with Welch’s correction, Figure 5D). Next, we analyzed the expression of postsynaptic density protein 95 (PSD95), a critical scaffolding protein for anchoring synaptic proteins including NMDA receptors (Lim et al., 2003; Won et al., 2016). No difference was observed in the neuropil fluorescent intensity along with the number of PSD95+ puncta between shRNA groups (p = 0.9919 and p = 0.1440, respectively; unpaired t-test with Welch’s correction, Figure 5B).

Additionally, we analyzed the effect of Ncan KO on vGLUT1+ and PSD95+ puncta (Figure 5E). Surprisingly, we did not find any significant differences in the fluorescence intensity and the number of vGLUT1+ puncta (p = 0.726 and p = 0.577, respectively; unpaired t-test with Welch’s correction, Figure 5F) and PSD95+ puncta (p = 0.826 and p = 0.9238, respectively; unpaired t-test with Welch’s correction, Figure 5F) between Ncan KO and wild-type mice.

### Effects of Ncan knockdown on activity-dependent ECM and synaptic remodeling

Finally, we studied the effects of Ncan knockdown on activity-induced alterations in the expression of ECM and synaptic plasticity markers. Therefore, we incubated mature cortical cultures (DIV21), infected with Ncan shRNA and Control shRNA at DIV7, in ASCF (basal condition) or ASCF supplemented with glycine to facilitate activation of NMDA receptors (GI-LTP condition) (Fortin et al., 2010), and quantified mRNA levels of selected candidate genes (Figure 6A). We opted for the glycine-induced form of chemical LTP (GI-LTP), the global form of LTP, as it has been shown to mediate similar cellular processes as tetanus-induced LTP (Chen et al., 2011). In Control shRNA-treated cultures, GI-LTP induced an increase in the basal mRNA expression of Ncan, Pcan, and TnC, (p = 0.046, p = 0.030 and p = 0.025, respectively; two-way ANOVA with Fisher’s LSD, Figure 6B) but not for Acan, Bcan, TnR and HAPLN1. However, in Ncan shRNA-infected cortical neurons, no such activity-dependent expression change was seen, except for HAPLN1 (p = 0.0007; two-way ANOVA with Fisher’s LSD, Figure 6B). Interestingly, GI-LTP upregulated HAPLN1 in the Ncan shRNA-treated group to similar levels seen in Control shRNA (1.182 ± 0.112 vs 1.193 ± 0.034; p = 0.939; two-way ANOVA with Fisher’s LSD, Figure 6B). A strong overall effect of shRNA on the mRNA expression levels of ECM was observed, except for HAPLN1 and TnR (Ncan: p < 0.0001; Acan: p < 0.0001; Bcan: p < 0.0001; Pcan: p = 0.0009; HAPLN1: p = 0.090, TnR: p = 0.895; TnC: p < 0.0001; two-way ANOVA with Fisher’s LSD, Figure 6B). On the other hand, GI-LTP only affected the expression of HAPLN1 compared to the other ECM molecules (Ncan: p = 0.241; Acan: p = 0.159; Bcan: p = 0.399; Pcan: p = 0.589; HAPLN1: p = 0.0012; TnC: p = 0.181; two-way ANOVA with Fisher’s LSD, Figure 6B). The interaction between shRNA and GI-LTP effects on the mRNA expression levels of ECM molecules was only seen for Pcan (p = 0.012; two-way ANOVA with Fisher’s LSD, Figure 6B).

**Figure 6.**
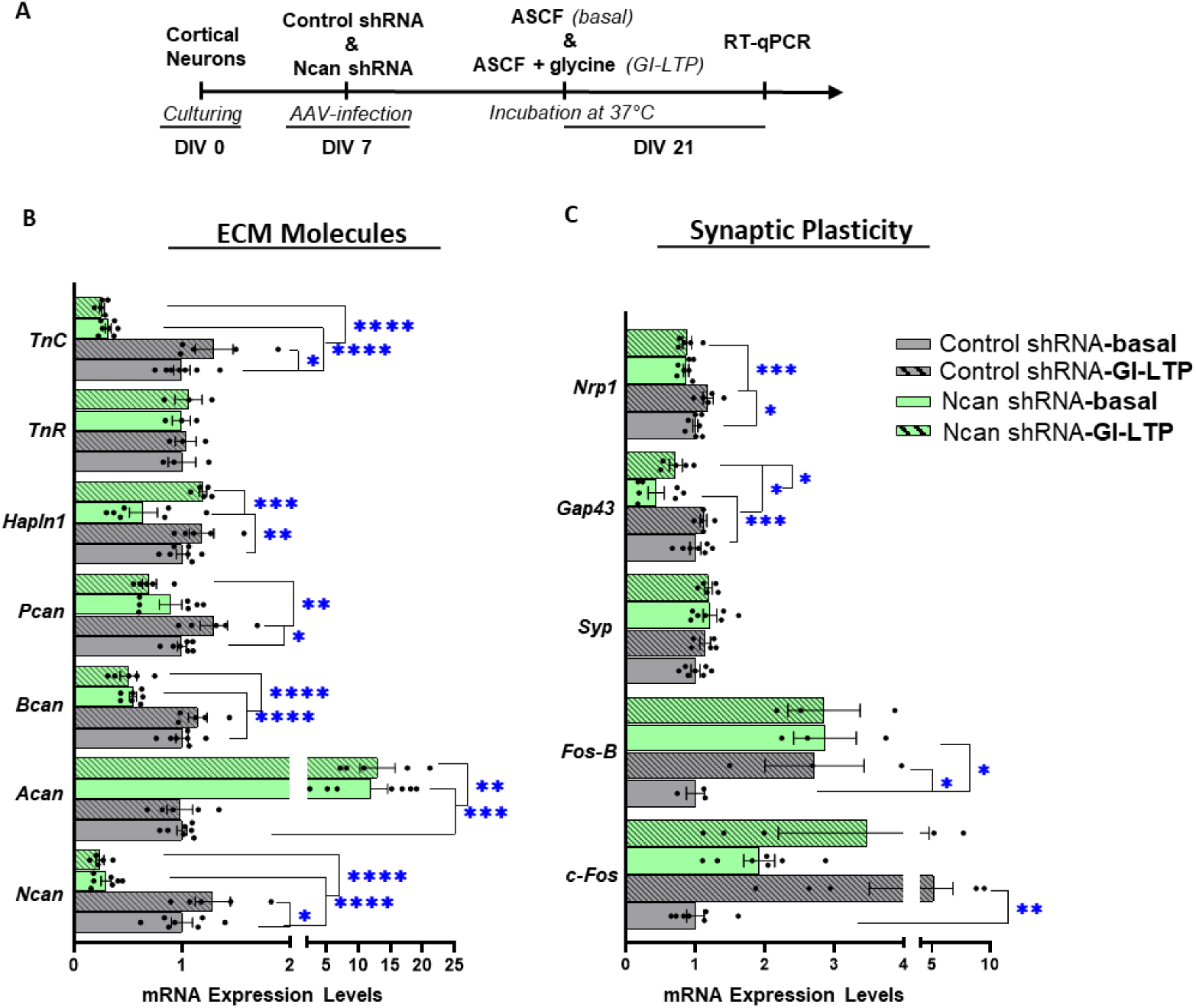
RT-qPCR analysis of basal and glycine-induced (GI-LTP) expression of ECM molecules and synaptic plasticity markers after Ncan knockdown in cultured cortical neurons. **(A)** Timeline showing the incubation of AAV-infected cortical neurons in ASCF (basal) and ASCF with 0.2 mM glycine (GI-LTP) at 37 ^0^C. The mRNA expression levels of ECM molecules **(B)** and synaptic plasticity markers **(C)** were determined. Bar graphs show mean ± SEM values. *p < 0.05, **p < 0.01, ***p < 0.001 and **** p < 0.0001 represent significant differences between Control shRNA (N=7 and 5 cultures in 2 independent experiments for basal and GI-LTP respectively) and Ncan shRNA (N=7 and 5 cultures in 2 independent experiments for both basal and GI-LTP) infected cortical neurons except for TnR and Fos-B (N=3 cultures in 2 independent experiments for basal and GI-LTP respectively) using two-way ANOVA with Fisher’s LSD.

Learning and memory change the expression of immediate-early genes such as c-Fos, whose expression are in turn critical for LTP maintenance and synaptic plasticity (Loebrich and Nedivi, 2009; Korb and Finkbeiner, 2011). Interestingly, LTP maintenance of stimulated CA1 pyramidal cells was reported to be disturbed in Ncan KO mice (Zhou et al., 2001). Therefore, we aimed to analyze the effect of Ncan knockdown on some synaptic plasticity markers under both basal and GI-LTP conditions. Activated by LTP-inducing neuronal activity, expression of c-Fos is involved in the regulation of transcription, and it is used as a neuronal activation marker (Gallo et al., 2018), while expression of Fos-B is considered to reflect basal neuronal activity (Nestler et al., 2001). Here, we show that there is a significant difference between the basal expression of Fos-B but not c-Fos in Ncan shRNA-relative to control shRNA-infected cortical neurons (p = 0.029 and p = 0.427, respectively; two-way ANOVA with Fisher’s LSD, Figure 6C).

Moreover, GI-LTP enhanced the basal expression of c-Fos and Fos-B in Control shRNA but not in Ncan shRNA (p = 0.003, p = 0.041, p = 0.224 and p = 0.985, respectively; two-way ANOVA with Fisher’s LSD, Figure 6C). These effects were specific as the expression of the most abundant presynaptic vesicular protein, synaptophysin (SYP), whose expression was shown to influence cognition as well as synaptic transmission (Tampellini, 2010; Xu et al., 2019), was not affected by Ncan knockdown.

Then, we checked other synaptic plasticity-associated targets such as the growth-associated protein 43 (GAP-43), which is an essential component of the presynaptic terminal remodeling in an activity-dependent manner (Benowitz and Routtenberg, 1997), and of axonal regeneration (Gil-Loyzaga et al., 2010). Interestingly, GAP-43 was downregulated in Ncan shRNA-infected cortical neurons under basal conditions compared to Control shRNA (p = 0.0002; two-way ANOVA with Fisher’s LSD, Figure 6C). GI-LTP enhanced the basal expression of GAP-43 in Ncan shRNA but not to control levels (p = 0.045 and p = 0.012, respectively; two-way ANOVA with Fisher’s LSD, Figure 6C). Additionally, the basal expression of neuropilin (NRP1), which regulates neuronal axon guidance and dendrite development (Tillo et al., 2012), was not altered between shRNA groups (p = 0.053; two-way ANOVA with Fisher’s LSD, Figure 6C). Strikingly, GI-LTP enhanced the expression of NRP1 in Control shRNA-but not in Ncan shRNA-infected cortical neurons (p = 0.019 and p = 0.840, respectively; two-way ANOVA with Fisher’s LSD, Figure 6C). This, therefore, resulted in a prominent difference in GI-LTP-dependent expression of NRP1 between shRNA groups (p = 0.0009; two-way ANOVA with Fisher’s LSD, Figure 6C). In summary, an overall significant effect of Ncan shRNA on the expression of synaptic markers was seen for GAP-43 and NRP1 (GAP-43: p < 0.0001; NRP1: p = 0.0003; two-way ANOVA with Fisher’s LSD, Figure 6C). These findings outline multiple molecular alterations in Ncan-depleted cortical neurons, potentially mediating the role of Ncan in synaptic plasticity.

## DISCUSSION

In this study, we provide evidence that Ncan is essential for PNN maintenance by influencing the mRNA expression levels of other PNN-associated molecules. Using shRNA against *Ncan* we were able to show that Ncan in the mPFC is critical for the temporal order recognition memory as well as for the consolidation/retrieval of spatial memories after reversal learning in the Labyrinth task. Moreover, we found that Ncan knockdown reduced perisomatic GABAergic innervation of PV+ interneurons and increased the density of vGLUT1+ presynapses in the neuropil of the PFC.

### Ncan knockdown and PNN structure

PNNs form and mature postnatally from around P14 to P100, and Ncan expression follows a similar trend (Pizzorusso et al., 2002). The composition of lecticans within PNNs has been shown to vary between brain regions (Lensjo et al., 2017; Fawcett et al., 2019). The expression of Acan, HAPLN1, TnR, and Pcan, but not Bcan, has recently been identified as essential contributors to PNN formation (Morawski et al., 2014; Suttkus et al., 2014; Eill et al., 2020). Additionally, a recent analysis of Ncan KO mice implicated Ncan as a major contributor to PNN integrity in the MNTB, where the authors reported alterations in the expression of other ECM molecules like decreased mRNA levels of HAPLN1 and Bcan (Schmidt et al., 2020). In the present study, we show that Ncan is a prominent factor for PNN maintenance. Our treatment with shRNA against Ncan resulted in about 25% reduction of Ncan immunoreactivity in PNNs. Although this reduction is below the reduction in the Ncan mRNA measured *in vitro*, yet it resulted in less prominent PNNs as was shown by WFA-staining that binds to the CSPGs’ glycosaminoglycan chains (Dityatev et al., 2007). Possible reasons for the rather small reduction of Ncan immunoreactivity can be expression of Ncan in non-infected neurons within mPFC or cells of different brain regions projecting to mPFC (Hoover and Vertes, 2007) or production and secretion of Ncan from astrocytes (Galtrey et al., 2008; Fawcett et al., 2019). We presume that astrocytic synthesis of Ncan was not affected in our study because the AAVs produced with the help of RapCap DJ largely do not target astrocytes *in vivo*, although it does *in vitro*. This makes the present data even more interesting by pointing to the role of neuronal Ncan in the PFC. Interestingly, we observed a linear relationship between reduced Ncan and WFA signals after Ncan knockdown. We also report a striking effect of Ncan knockdown on the mRNA expression levels of other PNN molecules such as downregulation of HAPLN1 and Bcan along with upregulation of Acan. This effect has also been reported in Ncan KO mice (Schmidt et al., 2020) and therefore supports the on-target efficacy of our knockdown vectors (Taxman et al., 2006; Okuda et al., 2014). Contrary to the KO study in the MNTB (Schmidt et al., 2020), the PNN structure was not strongly disrupted in the mPFC of Ncan-knockdown mice compared to wild-type, which may be either due to region-specific differences in PNN expression (Morikawa et al., 2017) or to Ncan reduction differences (100% vs. 25%). In summary, these data indicate neuronal Ncan as a prominent component and sculptor of PNNs around PV+ interneurons in mPFC.

### Neurocan depletion impairs mPFC-dependent cognitive functions

Neural ECM remodeling has been shown to impact cognition, both positively and negatively. For example, in TnR KO mice, ECM attenuation enhanced reversal learning in the Morris water maze and olfactory discrimination task (Morellini et al., 2010), as well as working memory (Sykova et al., 2005). Also, digestion of ECM with hyaluronidase improved reversal learning (Happel et al., 2014) and removal of ECM with chABC in the perirhinal cortex enhanced object recognition memory (Romberg et al., 2013). On the flip side, reduced LTP in hippocampal CA1 occurred in both TnR KO (Bukalo et al., 2001) and TnC KO mice (Evers et al., 2002) correlating with impaired fear extinction in the latter. Also, the Bcan KO impaired hippocampal-dependent LTP without obvious cognitive deficits (Brakebusch et al., 2002). However, the intra-hippocampal injection of hyaluronidase impaired LTP and contextual fear condition memory (Kochlamazashvili et al., 2010; Senkov et al., 2014). Additionally, digestion of ECM in the visual cortex disrupted recall of visual fear memory (Thompson and Chen, 2017) and more importantly in the mPFC, although in rats. Importantly, injection of chABC in the mPFC of rats resulted in a below chance-level performance in the cross-modal object recognition task (Paylor et al., 2018) and in an impairment of LTP (Shi et al., 2019). In the mPFC of anesthetized mice, chABC injection induced a decrease in gamma activity, a neuronal oscillation dependent on PV+ cells, which is essential for different cognitive tasks involving this neocortical region (Carceller et al., 2020).

In the present study, ECM attenuation through reduced Ncan expression impaired the temporal order recognition memory. Moreover, we found Ncan knockdown to negatively impact the retrieval of long-term reversal spatial memory (formed 22 ± 2 hours after the initial reversal learning phase) as well as the decision accuracy estimated in the percentage retrieval index approach. This method uses the mean number of errors, as studies have shown this measure to be the more effective metric compared to latency to reward (Maei et al., 2009) to measure cognition in labyrinth tasks (Rogers and Kesner, 2003; Lee and Kesner, 2004; Vago et al., 2007; Churchwell et al., 2010). Reduced Ncan expression in the mPFC did not influence spatial memory acquisition and recall during the learning phase within the same day but only during the memory tests on the next day. Our study is thus in line with a recent finding in humans highlighting similar functions of the mPFC, where different retrieval patterns were observed overnight relative to same-day memories (Ezzyat et al., 2018). This suggests that the ECM in the mPFC might be essential for the consolidation of updated spatial memory. One possible mechanism could involve the loss of the memory stabilization property of PNNs in Ncan-depleted mice (Gogolla et al., 2009; Leon et al., 2010). Also, the mPFC has been demonstrated to be critical for the retrieval of remote memories as well as consolidation and recall of recent memories (Leon et al., 2010) and these neural processes are PNN-dependent.

### Ncan reduction and increase in GABAergic inhibition

Changes in synaptic strength have been shown to contribute to learning and memory through various molecular mechanisms such as long-term depression (LTD) and LTP. LTP is produced when N-methyl-D-aspartate (NMDA) receptors and Ca^2+^ voltage-dependent channels are activated as a result of presynaptic release of neurotransmitters (e.g. glutamate) and postsynaptic depolarization (Mayford et al., 2012; Langille and Brown, 2018). This depends on the balanced interaction of excitation and inhibition (E/I balance) (Tao et al., 2014; Kirkwood, 2015), whereby variation in this balance affects basic cortical functions and may lead to impairment of mPFC-dependent learning (Yizhar et al., 2011). Moreover, the E/I balance is found to be critical in several psychiatric disorders such as autism and schizophrenia (Kirkwood, 2015). Interestingly, ECM modulation has been shown to affect the synaptic E/I balance in various model systems. For example, studies in TnR KO mice showed that deficiency in this ECM glycoprotein impairs the formation of perisomatic GABAergic synapses on pyramidal neurons, and as a result, decreased perisomatic inhibition in the CA1 region (Nikonenko et al., 2003; Saghatelyan et al., 2004) and enhanced the basal excitatory transmission, resulting in a shift in the E/I balance (Saghatelyan et al., 2004). Also, a strong effect of the ECM on the balance between inhibitory and excitatory synapses has been observed in quadruple KO mice, where the loss of TnR, TnC, Bcan, and Ncan increased the number of excitatory synapses and reduced inhibitory synapses (Gottschling et al., 2019). Interestingly, the expression of glutamic acid decarboxylase (GAD), the enzyme that synthesizes the inhibitory neurotransmitter, gamma-aminobutyric acid, depends on the ECM (Schmidt et al., 2020). As such, analysis of Ncan KO mice revealed a reduced expression of GAD 65/67 coupled with a prolonged delay of synaptic transmission, whereas the enzymatic removal of ECM decreased the density of cortical GABAergic perisomatic synapses (Sullivan et al., 2018; Schmidt et al., 2020). Other studies have found that PNN depletion induced a decrease in the density of puncta expressing inhibitory markers in the cortical neuropil or specifically in the perisomatic region of PV+ cells (Lensjo et al., 2017; Sullivan et al., 2018; Carceller et al., 2020). In line with these studies, we observed that both knockdown and knockout of Ncan significantly decreased the number and expression of vGAT puncta on PV+ interneurons. The underlying molecular mechanism dissected in *in vitro* studies may involve inhibition of NCAM-EphA3 association by Ncan’s interaction with the immunoglobulin-like domain 2 (Ig2) of NCAM (Sullivan et al., 2018).

However, knockdown of Ncan also upregulated vGLUT1 (although not vGLUT2) expression. A previous study revealed that Ncan inhibited Semaphorin 3 (SEMA3)-dependent spine pruning (which happens mostly at excitatory postsynaptic sites), and consequently, Ncan depletion could have the opposite effect, thereby increasing the number of spines and excitatory synapses (Mohan et al., 2018). Additionally, previous studies have also shown that the expression of vGLUT2 is independent of the upregulation in vGLUT1 (Wojcik et al., 2004), in line with a specific effect of Ncan knockdown on vGLUT1+ presynapses. Overall, these data demonstrate that Ncan knockdown leads to impaired connectivity within the PV+ cell network and modulation of excitatory input to principal cortical cells, which may affect PFC network activity in a complex manner, requiring further functional characterization of cells and the network, to conclude that Ncan depletion may change the E/I balance in the PFC in favor of excitation.

### Ncan knockdown and synaptic plasticity

To further study the effects of Ncan downregulation on mPFC-dependent cognitive processes, we searched for expression changes of multiple PNN components and synaptic plasticity-related genes under basal vs. GI-LTP conditions. One major finding was the reduced expression of GAP-43 in both basal and GI-LTP conditions. Previous studies have shown a strong relationship between GAP-43 expression and memory. For example, overexpression of GAP-43, as seen in the hippocampus of Alzheimer’s patients, impaired learning and memory whereas moderate expression improved it (Rekart et al., 2005; Holahan et al., 2007). Reduced expression of GAP-43 resulted in memory dysfunction similar to the effect seen in the present study (Rekart et al., 2005).

On the other hand, the enhanced basal expression of Fos-B observed after Ncan knockdown in cortical neurons is also an interesting finding as its expression strongly correlates with neural epileptiform activity (Vossel et al., 2016), in line with our immunohistochemical data showing impaired GABAergic inhibition of PV+ interneurons and increased numbers of vGLUT1+ puncta in Ncan knockdown. In AD patients and several AD animal models, subclinical epileptiform activity has been reported, which correlates strongly with the chronic accumulation of Fos-B, impacting also the expression of c-Fos (Kam et al., 2016; Corbett et al., 2017). Interestingly, overexpression of Fos-B is also shown to impact hippocampal learning and memory, possibly through Fos-B dependent induction of immature spines (Eagle et al., 2015), which might be related to the cognitive deficits seen in Ncan shRNA mice. Moreover, the acute neuronal activity-dependent upregulation of c-Fos that is essential for LTP-dependent synaptic plasticity (Calais et al., 2013), i.e. the cellular and molecular form of learning and memory (Flavell and Greenberg, 2008), did not occur in cortical cells after Ncan knockdown. This is in agreement with a previous work in which basal c-Fos expression was found to be upregulated in memory-impaired aged mice (Haberman et al., 2017). Interestingly, NRP1 expression was also affected by Ncan knockdown, and since it is a receptor for SEMA3A (Raimondi et al., 2016) during axonal pathfinding, dendrite development and branching (Tillo et al., 2012), it is possible that Ncan might modulate signaling through the SEMA3-NRP1 pathway (Fenstermaker et al., 2004).

In conclusion, our study highlights the importance of PNNs and perisynaptic ECM associated with excitatory vGLUT1+ synapses for synaptic wiring in the PFC, temporal order recognition memory as well as the consolidation/retrieval of updated spatial memories. Overall, our results suggest that the ECM composition and integrity within the mPFC confer critical modulatory effects on cognitive flexibility.

## Acknowledgments

We thank Katrin Boehm and Jenny Schneeberg for their technical support. This work has been supported by DAAD (fellowship to D.B.A), Deutsche Forschungsgemeinschaft (DFG, German Research Foundation, projects 425899996 / SFB 1436 and 362321501/RTG 2413 SynAGE to A.D. and C.S.) and Spanish Ministry of Science and Innovation (RTI2018-098269-B-I00 to J.N.) and the Generalitat Valenciana (PROMETEU/2020/024 to J.N.).

## Author’s contribution

D.B.A. performed all behavior experiments and with L.S did IHC analyses after Ncan knockdown; H.M performed all RT-qPCR analyses, H.C., M.P.R. along with B.G.V. performed all experiments in Ncan-KO mice; A.D. with J.N and R.K designed the study and supervised data analysis. D.B.A. wrote a draft of the manuscript and A.D., C.S. and J.N. edited the manuscript; C.S. supervised breeding of Ncan KO mice; R.K designed Ncan shRNA AAVs and has written Fiji scripts for image analysis of all samples derived from shRNA-treated mice, which were modified by D.B.A..

---

**Table 1.**
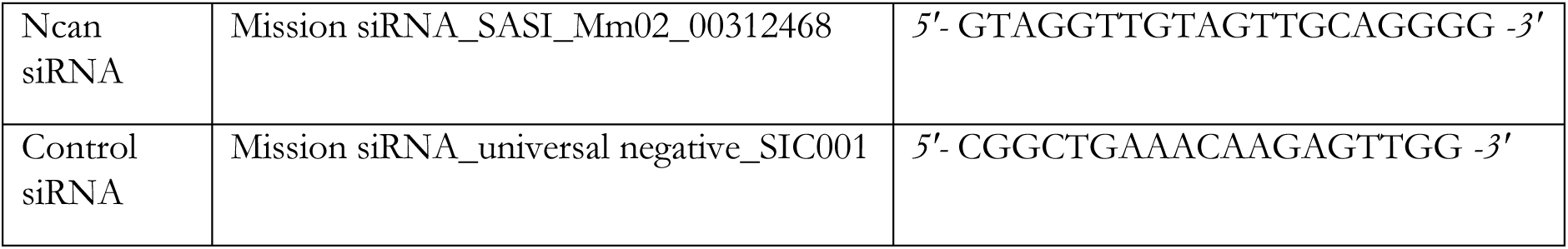
SiRNA sequences used in this study.

