## Supplementary tables and figures for "Depletion of neurocan in the prefrontal cortex impairs temporal order recognition, cognitive flexibility and perisomatic GABAergic innervation": Ncan suppl matter.pdf

**Supplementary Table 1. Full Description of TaqMan™ probes used for qPCR**

| Protein | Gene | Species | Reference Sequence | | Sequence (5' to 3') |
|---|---|---|---|---|---|
| ACAN | *Acan* | *Mus musculus* | NM_007424.2 | Fw | CTTACCCTGAGGCTGGTGTG |
| | | | | Rv | ACATTGCTCCTGGTCTGCAA |
| Ncan | *Ncan* | *Mus musculus* | NM_007789.3 | Fw | CAGGCCACAGCAATCATCCT |
| | | | | Rv | GATGTGGAAACAGAAGTGGGG |
| BCAN | *Bcan* | *Mus musculus* | NM_001109758.1 | Fw | TGCCCTCGTTCCCTTTTCTG |
| | | | | Rv | CTGTGTGGCCAGTGAGATGT |
| PCAN | *Pcan* | *Mus musculus* | NM_001081306.1 | Fw | AACAGTTCAGAGGCAGAGGC |
| | | | | Rv | CAGCTCTGCACTTCCTGGTA |
| HAPLN1 | *Hapln1* | *Mus musculus* | NM_013500.4 | Fw | AGTCTCCTGGTGACGCTTTG |
| | | | | Rv | GGGGCCATTTTCTGCTTGGA |
| TnR | *TnR* | *Mus musculus* | NM_022312.3 | Fw | AGTAAGAGGGGAAAAGAGAG |
| | | | | Rv | TCGTAAATCA TTGTCGTGTA |
| TnC | *TnC* | *Mus musculus* | NM_011607.3 | Fw | GACACCTAGCCAGTCCAACC |
| | | | | Rv | GCAGACACACTCGTTCTCCA |
| GAPDH | *Gapdh* | *Mus musculus* | NM_001289726.1 | Fw | CCCTTAAGAGGGATGCTGCC |
| | | | | Rv | ACTGTGCCGTTGAATTTGCC |
| c-FOS | *c-fos* | *Mus musculus* | NM_010234.2 | Fw | TGATGTTCTCGGGTTTCAAC |
| | | | | Rv | ACCCACGCTGCTGGCCCTGT |
| ARC | *Arc* | *Mus musculus* | NM_001276684.1 | Fw | GGAGCTGGACCATATGACCA |
| | | | | Rv | CAGGACCCAGCCTGAATAGA |
| GAP43 | *Gap43* | *Mus musculus* | NM_008083.2 | Fw | ATGCTGTGCTG TATGAGAAG |
| | | | | Rv | AGGCTGACCAAGAACATGCC |
| FOS-B | *Fos-B* | *Mus musculus* | NM_008036.2 | Fw | TGCATCGAAACTTGGGCAGT |
| | | | | Rv | AAACCCGCAAGGAACAAGGA |
| NRP1 | *Nrp1* | *Mus musculus* | NM_008737.2 | Fw | TTCATTGCTCTCTCTCCTTC |
| | | | | Rv | CGTTTTCATAAAAATCCCAAA |
| SYP | *Syp* | *Mus musculus* | NM_009305.2 | Fw | GGTGCAAGGGGCGTGGCTGC |
| | | | | Rv | TAAACTCCTCAACTATCTAGTC |

**Supplementary Table 2. Full Description Primary Antibody**

| Antibody | Host | Isotype | Dilution | Company | Antibody ID |
|---|---|---|---|---|---|
| PV | Mouse | Monoclonal | 1:500 | Sigma-Aldrich | P3088 |
| PV | Chicken | Polyclonal | 1:500 | Abcam | AB2619887 |
| WFA | Streptavidin | Biotinylated | 1:1000 | BioLogo (Vector Laboratories) | B1355 |
| vGAT | Guinea pig | Polyclonal | 1:500 | Abcam | AB887873 |
| vGAT | Rabbit | Polyclonal | 1:500 | Synaptic systems | 131002 |
| PSD 95 | Mouse | Monoclonal | 1:250 | Neuromab | 73-028 |
| vGLUT1 | Guinea Pig | Polyclonal | 1:2000 | Synaptic systems | 135304 |
| Ncan | Sheep | Polyclonal | 1:100 | R&D Systems | AF5800 |
| vGLUT2 | Mouse | Monoclonal | 1:500 | Abcam | AB79157 |
| CAMKII | Mouse | Monoclonal | 1;500 | Abcam | AB22609 |

**Supplementary Table 3. Full Description Secondary Antibody.**

| Anti | Host | Label | Dilution | Company | Antibody ID |
|---|---|---|---|---|---|
| Chicken | Goat | A647 | 1:400 | Life Technologies (Invitrogen) | A21449 |
| Guinea Pig | Goat | A555 | 1:400 | Life Technologies (Invitrogen) | A11073 |
| Mouse | Goat | A488 | 1:400 | Life Technologies (Invitrogen) | A11029 |
| Mouse | Goat | A405 | 1:1000 | Life Technologies (Invitrogen) | A31553 |
| Streptavidin | Goat | A647 | 1:1000 | Life Technologies (Invitrogen) | S21374 |
| Rabbit | Goat | A546 | 1:1000 | Life Technologies (Invitrogen) | A11035 |
| Mouse | Goat | A546 | 1:1000 | Life Technologies (Invitrogen) | A11030 |
| Guinea Pig | Goat | A647 | 1:1000 | Life Technologies (Invitrogen) | A21236 |
| Sheep | Donkey | A546 | 1:1000 | Life Technologies (Invitrogen) | A21098 |
| Mouse | Goat | A546 | 1:1000 | Life Technologies (Invitrogen) | A11030 |

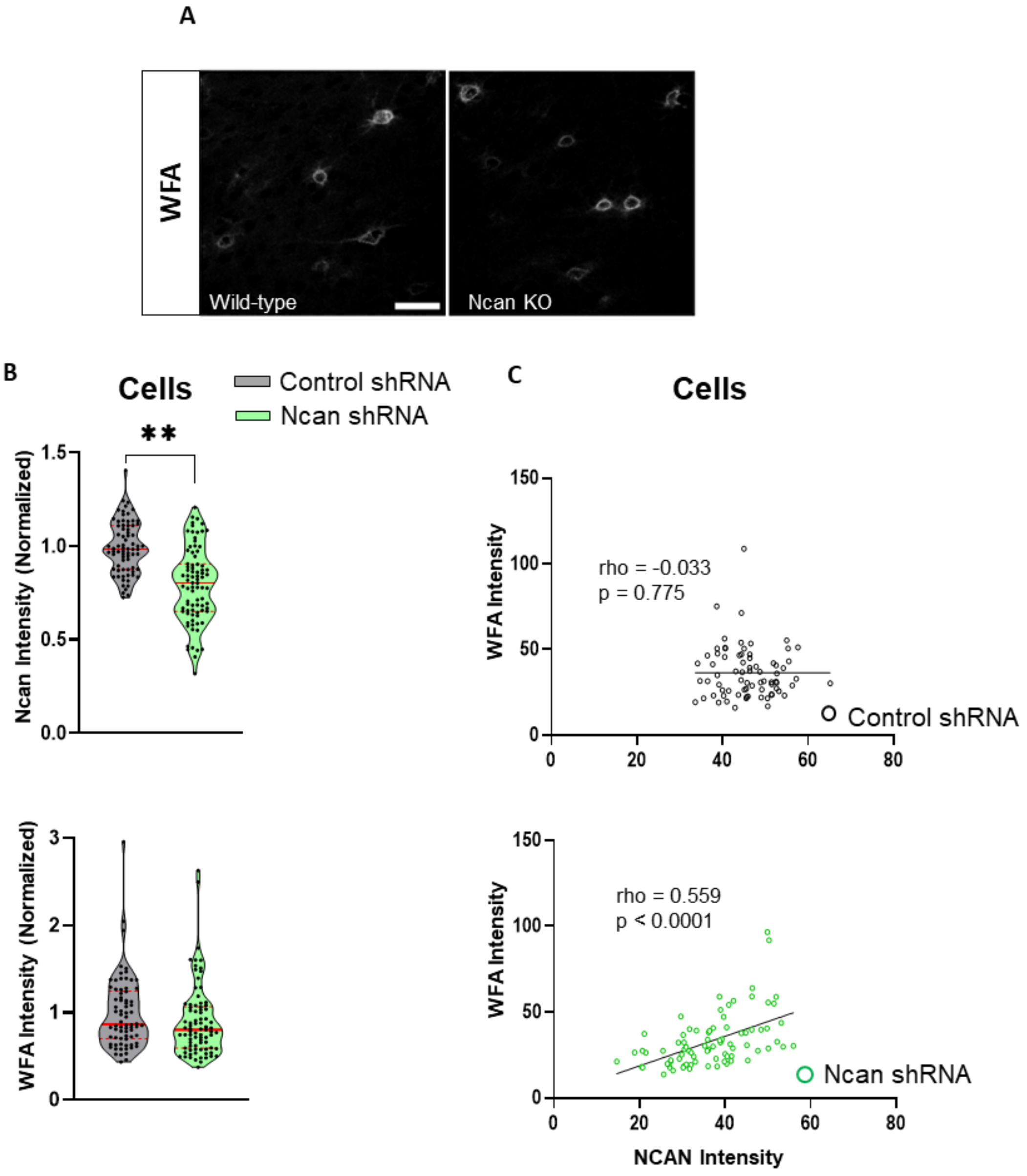


**Figure S1. Knockdown of Ncan expression in the PFC.**

**(A)** Representative images of PNNs that were labelled with WFA visualized after Ncan KO. Scale bar, 40 μm. **(B)** The perisomatic expression of Ncan per cell was reduced in Ncan shRNA AAV-injected mice compared to Control shRNA AAV-injected mice., but not for WFA. **(C)** Additionally, a positive correlation between the expression of Ncan and WFA in Ncan shRNA mice but not for Control shRNA**.** Bar graphs show mean ± SEM values. *$p < 0.05$, **$p < 0.01$, ***$p < 0.001$ and **** $p < 0.0001$, represent significant differences between wild-type (N = 5) and Ncan KO (N = 5) mice or Control shRNA (N = 7) and Ncan shRNA (N = 9) treated mice using one-way ANOVA with Sidak's multiple comparisons test; Brown-Forsythe and Welch ANOVA tests; nested t-test after log transformation of WFA intensities and no transformation of Ncan intensities; unpaired t-test with Welch's correction, or significant Spearman coefficients of correlation.

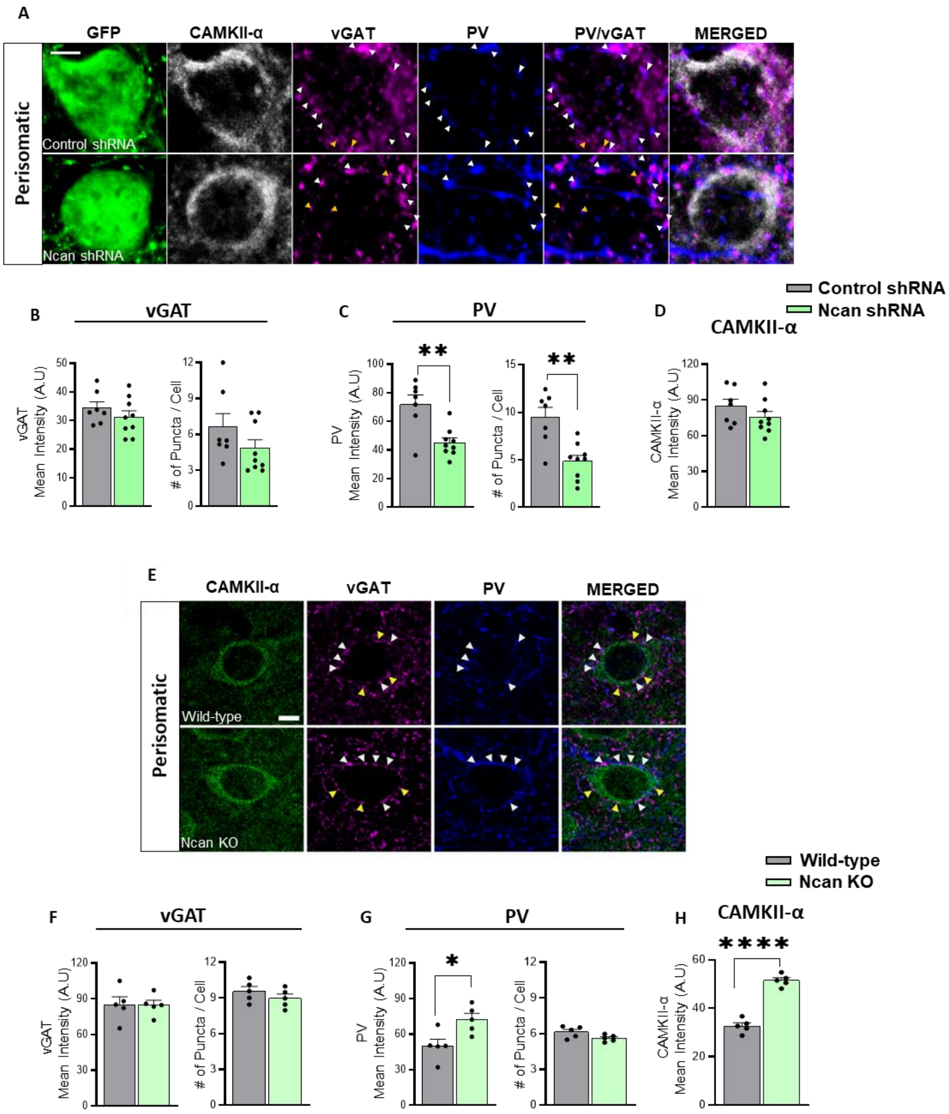


**Figure S2. Effects of Ncan knockout on GABAergic innervation of excitatory pyramidal neurons.**

The perisomatic expression and the number of vGAT and PV puncta were analyzed in Ncan-deficient mice as well as in Ncan shRNA mice compared to their respective controls. **(A)** Representative images showing vGAT+ and PV+ puncta contacting the somata of CaMKII-α expressing pyramidal neurons in shRNA-treated mice. Scale bar, 5 µm. White arrowheads point to the puncta co-expressing both PV and vGAT, whereas yellow arrowheads point to those puncta expressing VGAT but not PV. **(B)** No difference was observed for the expression and number of vGAT puncta in Ncan shRNA and Control shRNA mice. **(C)** However, the expression and number of PV+ puncta were decreased in Ncan shRNA mice. **(D)** No difference in the fluorescent intensity of CAMKII-α was detected. **(E)** Representative images showing vGAT+ and PV+ puncta contacting the somata of CaMKII-α expressing pyramidal neurons in Ncan KO and wild-type mice. Scale bar, 5 µm. **(F)** No difference in the expression and number of vGAT+ puncta between Ncan KO and wild-type mice was observed. **(G)** However, Ncan KO upregulated the PV expression but not the number of PV puncta compared to wild-type mice. **(H)** Interestingly, somatic CAMKII-α expression was upregulated in Ncan KO mice compared to wild-type mice. Scale bar, 5 µm. Bar graphs show mean ± SEM values. *$p < 0.05$, **$p < 0.01$, ***$p < 0.001$ and **** $p < 0.0001$ represent significant differences between wild-type ($N = 5$) and Ncan KO ($N = 5$) mice or Control shRNA ($N = 7$) and Ncan shRNA ($N = 9$) using an unpaired t-test with Welch's correction.
